# Computational Framework to Predict and Shape Human-Machine Interactions in Closed-loop, Co-adaptive Neural Interfaces

**DOI:** 10.1101/2024.05.23.595598

**Authors:** Maneeshika M. Madduri, Momona Yamagami, Si Jia Li, Sasha Burckhardt, Samuel A. Burden, Amy L. Orsborn

## Abstract

Neural interfaces can restore or augment human sensorimotor capabilities by converting high-bandwidth biological signals into control signals for an external device via a decoder algorithm. Leveraging user and decoder adaptation to create co-adaptive interfaces presents opportunities to improve usability and personalize devices. However, we lack principled methods to model and optimize the complex two-learner dynamics that arise in co-adaptive interfaces. Here, we present computational methods based on control theory and game theory to analyze and generate predictions for user-decoder co-adaptive outcomes in continuous interactions. We tested these computational methods using an experimental platform where human participants (N=14) learn to control a cursor using an adaptive myoelectric interface to track a target on a computer display. Our framework allowed us to characterize user and decoder changes within co-adaptive myoelectric interfaces. Our framework further allowed us to predict how decoder algorithm changes impacted co-adaptive interface performance and revealed how interface properties can shape user behavior. Our findings demonstrate an experimentally-validated computational framework that can be used to design user-decoder interactions in closed-loop, co-adaptive neural interfaces. This framework opens future opportunities to optimize co-adaptive neural interfaces to expand the performance and application domains for neural interfaces.

## 1 Introduction

Neural interfaces can restore or augment human capabilities [1–6]. In a neural interface, signals from the user are translated via a decoder algorithm to control a device. Decoder algorithms that adapt to users during interface operation can improve performance and enable personalization [2, 7–17]. Users also adapt as they learn to control the interface because they receive real-time feedback [3, 18–21]. Introducing adaptive algorithms into neural interfaces therefore creates a co-adaptive system. Prior work has demonstrated the benefits of leveraging co-adaptation to guide or assist user adaptation [10, 22, 23] and maintain performance over time [10, 14, 24].

However, co-adaptive systems are challenging to design because they involve dynamic, two-learner interactions: both the user and decoder may adapt simultaneously and in response to one another. Experiments have exposed the importance of identifying appropriate decoder parameters, such as the rate of decoder adaptation, to ensure stable and predictable interface control [7, 25, 26]. Methods to design and analyze two-learner systems are an active area of research in neural engineering [27, 28] and dynamic game theory [29], but co-adaptive neural interfaces are largely designed and implemented empirically. Theoretically-grounded and experimentally-validated techniques to model and quantify co-adaptive dynamics will unlock new ways to design robust interfaces that harness the full potential of user and decoder adaptation.

Existing frameworks for designing co-adaptive interfaces highlight the promise of model-based approaches but either do not capture the full range of dynamic interactions that can arise between users and decoders, or do not make predictions about interaction outcomes. Past work on neural interfaces focused on the user adapting to a leading, optimal (fixed) decoder [30] or the decoder adapting to follow the user [31], and most experimental neural interfaces are constrained to this limited range of co-adaptive interactions [2, 10, 14, 22]. We instead seek methods that will be able to accommodate a wide spectrum of user-decoder interactions, where the device may shift from ‘leading’ to ‘following’ the user over time [32]. Game theory has been used to analyze the joint responses of two learners in human-human motor interactions [9, 33] and human-machine interactions [25, 34]. We built on these ideas, adapting tools from control theory and game theory to create a flexible, experimentally-validated framework for analysis and synthesis of co-adaptive outcomes of continuous interactions in neural interfaces.

We developed a myoelectric interface platform that allowed us to build experimentally-validated frameworks for co-adaptive neural interfaces. We validated that users changed their behavior alongside an adaptive decoder to control a cursor via their muscle activity, creating a co-adaptive system. We modeled this closed-loop system using control theory methods to quantify interactions between users and the device. This experimental platform and analysis toolkit allowed us to test analytical predictions about user-machine behaviors derived from a model treating the user and decoder as two agents in a game. Our experimental results revealed that user learning can be influenced through real-time interface adaptation and that these outcomes can be predicted using game-theoretic models. Together, our findings establish methods to analyze and synthesize co-adaptive systems that can enable next-generation approaches for designing and personalizing neural interfaces.

## 2 Results

We created an experimental platform to study co-adaptation using non-invasive myoelectric interfaces, which we subsequently used as a testbed to develop and validate computational frameworks for coadaptive interactions. This platform created unfamiliar human-machine interactions for our human participants (N=14) which let us systematically manipulate decoder properties to investigate the effect of decoder adaptation on user adaptation. Here, we use the term “adaptation” to describe *any change over time* in the decoder or user, and do not aim to draw analogies to particular forms of motor learning [35]. Our myoelectric interface had similar properties to many common motor neural interfaces, including brain-computer interfaces: it used high-dimensional biosignals to control a lower-dimensional movement [36, 37] and was a closed-loop system that can facilitate user adaptation [12]. We first analyzed the experimental platform to establish that both users and decoders adapt, creating a co-adaptive system (section 2.1). We then leveraged control theory and game theory to develop computational methods that can measure and predict co-adaptive outcomes (sections 2.2 and 2.3, respectively). Finally, we used our experimental platform to validate model predictions by manipulating the decoder adaptation parameters and demonstrating that our framework could predict outcomes of co-adaptive interactions (sections 2.4 and 2.5).

### 2.1 An adaptive myoelectric interface created a co-adaptive experimental platform

Participants completed a 2D trajectory-tracking task using a 64-channel adaptive myoelectric interface (Fig. 1A, Extended Data Fig. 1A). The task was chosen for its advantages in analyzing user sensorimotor transforms [38, 39]. Participants controlled a cursor using surface electromyography (sEMG) signals measured from their dominant forearm via a velocity-controlled Wiener filter decoder (see Methods). The decoder was randomly initialized at the beginning of each trial and then updated online every 20 seconds using previously-developed adaptive decoding algorithms [40, SmootBatch]. Decoder adaptation conditions varied from trial to trial, and participants completed a total of 16 5-minute trials (arranged in 2 blocks of 8 trials) (Fig. 1B, see Methods).

**Fig. 1.**
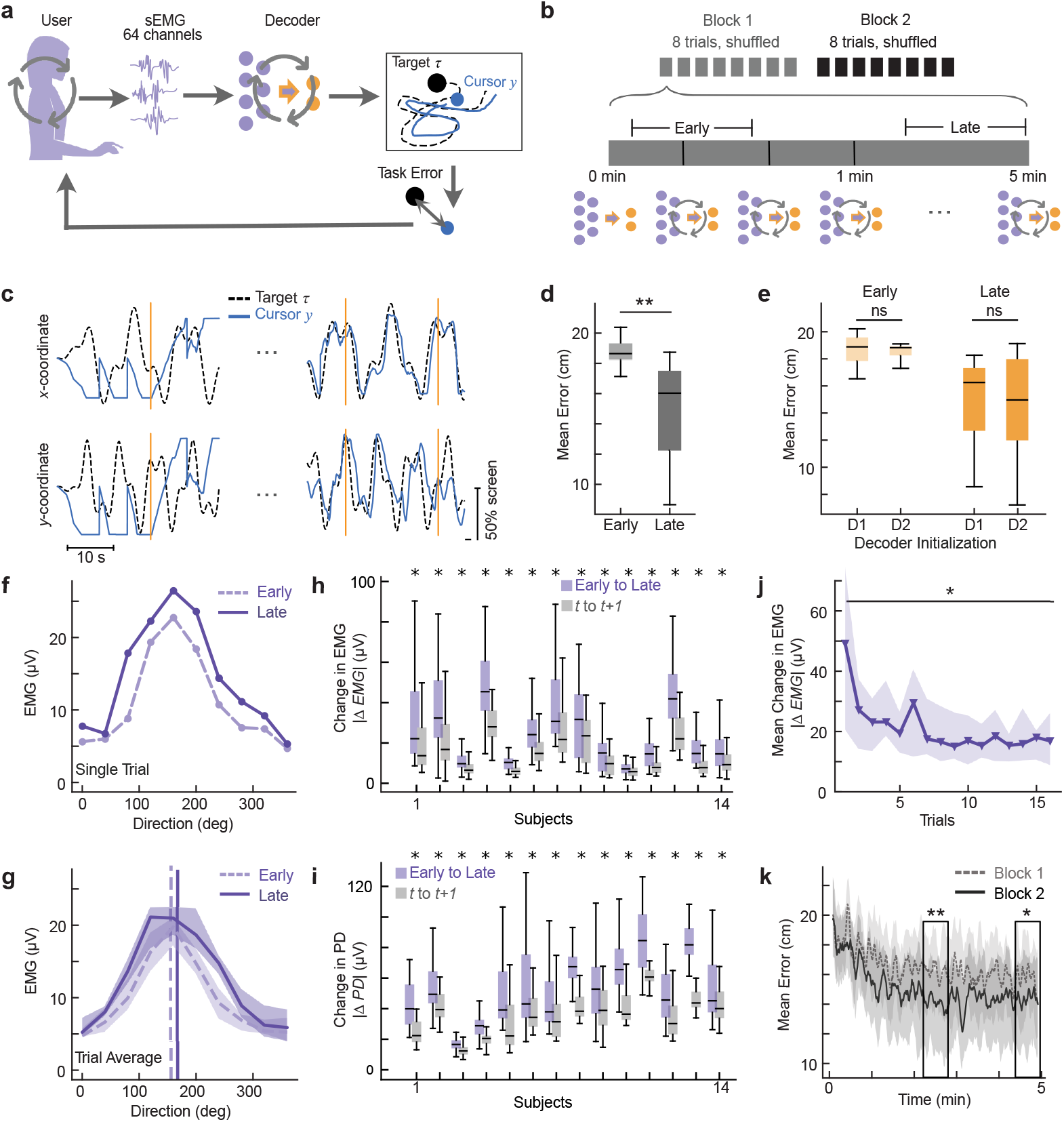
Co-adaptative myoelectric interface. **a**. Experiment schematic. Users tracked a target by controlling 2D cursor velocity with muscle activity via an adaptive decoder. **b**. Session schematic. Participants completed two blocks of eight 5-minute trials with different, randomized, decoder conditions; decoders were initialized randomly and adapted every 20 seconds. **c**. Example cursor (solid blue) and target (dashed black) trajectories. Orange lines denote decoder updates. ^1^**d**. Mean error (N = 14, center shows median; box shows 25th–75th percentiles; whiskers extend to 1.5 × interquartile range) early (first 30 s) and late (last 30 s) in trials. Error (∥*τ* −*y*∥) was averaged across trials for each participant. One-sided Wilcoxon signed-rank test, **p = 0.0001. **e**. Mean error (N = 14) separated by decoder initialization (D1, D2) early and late in trials. Error (∥*τ* −*y*∥) was averaged across trials for each participant. Two-sided Wilcoxon signed-rank test, (Early) ns = 0.67, (Late) ns = 0.43. **f**. EMG tuning curve for one EMG channel early (dashed light purple) and late (solid purple) within a single trial. **g**. Average EMG tuning curve for one channel early (dashed light purple) and late (solid purple) across all trials (N = 16, median; shading shows the 25th - 75th percentile range). Preferred directions for each curve shown with vertical lines. **h**. Norm difference of the average early versus late EMG tuning curves (∥Δ*EMG*∥ = ∥*EMG*(*late*) −*EMG*(*early*)∥) (purple) compared to norm difference of the average consecutive 30-second intervals (∥Δ*EMG* ∥= ∥*EMG*(*t*) −*EMG*(*t* + 1) ∥) (gray) for all channels, computed for each subject (N = 63, One-sided Wilcoxon signed-rank test with a Bonferroni correction of n = 14; *p *<* 0.001 for all subjects). **i**. Magnitude of angular change in EMG preferred direction early versus late (∥ Δ*PD*∥ = ∥*PD*(*late*) −*PD*(*early*) ∥) (purple) compared to angular change in consecutive 30-second intervals (∥Δ*PD*∥ = ∥*PD*(*t*) −*PD*(*t* −1) ∥) (gray)for all channels, computed for each subject (N = 63, One-sided Wilcoxon signed-rank test with a Bonferroni correction; *p *<* 0.001 for all subjects). **J**. Average change in EMG tuning curves within a trial (∥Δ*EMG*∥ = ∥*EMG*(*late*) −*EMG*(*early*) ∥) for all subjects as a function of trial number (N = 14, median; shading shows 25th - 75th percentile range). Average change in EMG of first trial is compared to last trial. (One-sided Wilcoxon signed-rank test, *p = 0.0083). **K**. Average error as a function of trial time in block 1 (dashed gray) and block 2 (solid black) (N = 14, median; shading shows 25th-75th percentile range). Black boxes represent comparisons of mid-trial and end-trial task error between block 1 and block 2 (One-sided Wilcoxon signed-rank test, **p = 0.00043, *p = 0.015). All boxplots (panels D, E, H, and I) show the median (center line), 25th-75th percentiles (box; interquartile range), and 1.5 x interquartile range (whiskers).

We first analyze general properties of system behavior independent of decoder adaptation conditions before examining the influence of decoder conditions (learning rate and cost functions, sections 2.4 and 2.5, respectively). Performance improved as the decoder adapted during a trial (Wilcoxon signed-rank test, *p <* 0.001) (Fig. 1C, D) and these performance improvements were not impacted by decoder initialization (Wilcoxon signed-rank test, *p >* 0.05) (Fig. 1E), similar to invasive brain-computer interface adaptive decoder studies [27, 40, 41]. These results confirm expected behavior of the adaptive algorithms.

While past work suggests that users likely adapt alongside the algorithm, we next aimed to confirm this behavior in our testbed. System-level performance metrics alone cannot separate the effect of user adaptation from that of decoder adaptation because performance changes could be attributed to either learner. The decoder adapted during the trial by design, leading to shifts in the decoder matrix (*D*) according to prescribed cost functions (e.g., Fig. 5C, also reported in preliminary studies in 26). To validate that our myoelectric interfaces created a co-adaptive system, we examined whether users adapted during each trial. We asked whether the user’s behavior changed while controlling the interface by estimating the relationship between user sEMG activity and the direction of intended movement – i.e., an EMG tuning curve [42–44] (see Methods). Variation in electrode placement and user behavior resulted in a wide range of EMG tuning curves that varied across participants and across electrodes (Extended Data Fig. 2). EMG tuning curves from a sample participant revealed within-trial changes in user behavior (Fig. 1F, G). We saw a range of changes in EMG tuning curves in other participants (Extended Data Fig. 3). We quantified EMG changes for each participant by calculating the magnitude of the difference between EMG tuning curves (Fig. 1H) and angular differences in the preferred direction (Fig. 1I) (see Methods). Because changes in EMG tuning were diverse across electrodes and participants, we assayed whether changes across a trial reflected a directed process. We compared the magnitude of change in the first 30 seconds (early) versus the last 30 seconds (late) to the magnitude of change on consecutive 30 second intervals. All participants showed significantly larger changes in the magnitude and tuning angles of EMG activity over the course of the trial than moment-to-moment fluctuations, suggesting that users adapted their behavior in a directed fashion during the 5-minute trial (Wilcoxon signed-rank test with a Bonferroni correction of *n* = 14, *p <* 0.001) (Fig. 1H, I).

Interestingly, we observed that within-trial differences in EMG tuning curves decreased as trials progressed (Fig. 1J). We hypothesized that these longer timescale changes reflected users learning over the course of the experiment. We tested this prediction by comparing performance in the first and second blocks, which contained trials with matched decoder adaptation conditions. Decoder adaptation parameters were matched across blocks of trials, which means that differences in task performance (determined by the joint user-decoder system) must stem from differences in user behavior between blocks. We indeed saw that participants’ average mid-trial and final task error was lower in block 2 compared to block 1 (Wilcoxon signed-rank test, *p <* 0.05), suggesting that task performance trajectories improved more and more quickly in block 2 than in block 1 (Fig. 1K). Taken together, user EMG and system performance suggest that users adapt alongside the decoder in our myoelectric interface, both within and across trials, establishing a co-adaptive system.

### 2.2 Co-adaptation outcomes matched control theory predictions

Distinguishing user and decoder contributions to an interface is key to understanding and designing closed-loop, co-adaptive systems. While decoders are known exactly, the user model must be estimated. The EMG direction-tuning analysis we used above can characterize user activity but does not provide a model for the user and, therefore, limits our ability to examine user-decoder interactions. We used methods from control theory, a discipline focused on analysis and synthesis of feedback systems, to model the behavior of users and the closed-loop system. Prior work shows that control theory can estimate changes in user trajectory-tracking behavior by separating their feedback and feedforward controllers [38, 45–47]. We extended these techniques to our multi-input, multi-output system to analyze co-adaptive outcomes. These approaches involve modeling users as performing position-based control with perfect and immediate sensory feedback. These models, therefore, notably simplify computations known to contribute to motor control (such as sensory delays). Yet, these and related simplified models have proven useful for capturing aspects of motor behavior in humans and other model organisms [39, 48, 49].

We define our model of the user’s encoder as the closed-loop mapping between task information and EMG activity in our myoelectric interface (Fig. 2A; see Extended Data Table 1 for a summary of variables). Our encoder model, *E* ∈ℝ^64×8^, considers feedforward and feedback inputs to the user, which are derived from closed-loop task information presented to the user - target position (*τ* ∈ ℝ^2^), target velocity 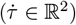, position error (*τ* ∈*y* −ℝ^2^) and velocity error 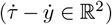. We formulate an encoder model with these feedforward and feedback elements that depend linearly on target position, cursor error, and an offset *β* (to capture resting activity):

**Table 1.**
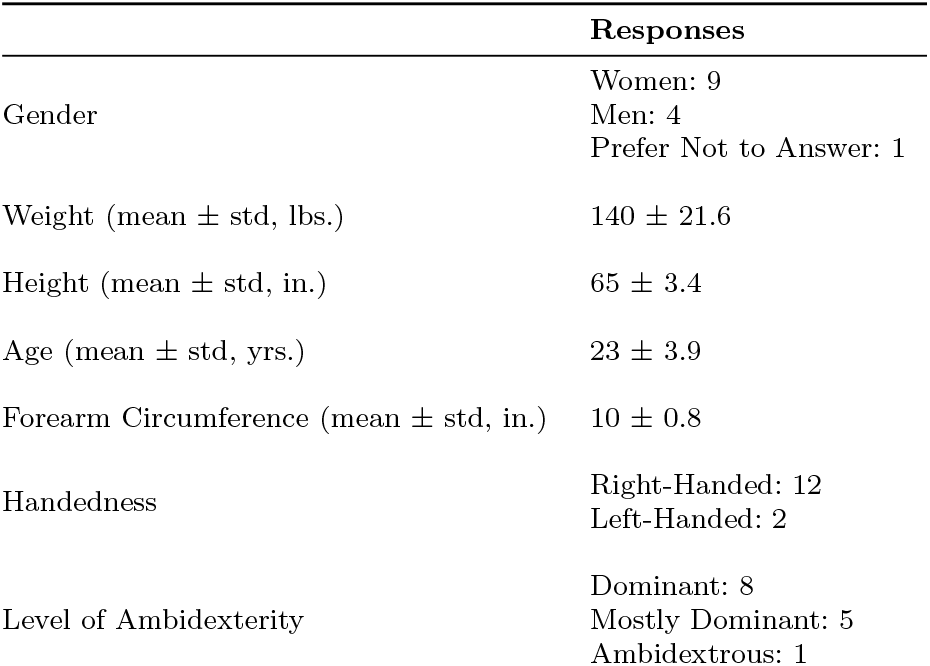
Participant Demographics.

**Fig. 2.**
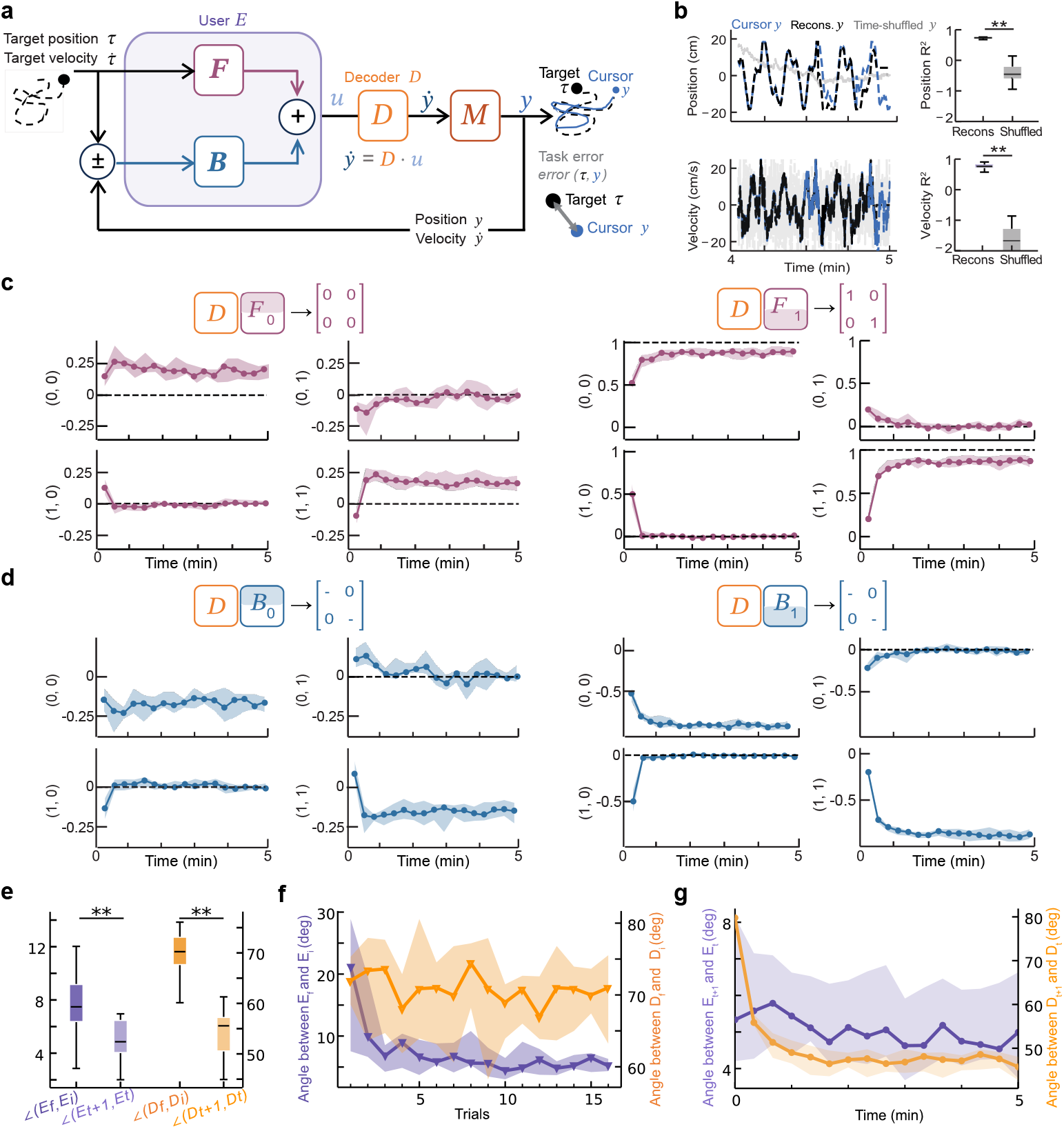
Control-theoretic analysis allows for user encoder estimation and decoder-encoder analysis. **a**. Block diagram model of closed-loop system. The user’s encoder *E* outputs myoelectric activity *u* to the decoder *D* to follow the target’s position and velocity. The decoder outputs velocity, which is then integrated by system dynamics *M* to generate a cursor position. The user sees their cursor moving with some velocity 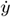 to position *y*. Error in the cursor position and velocity (relative to the target) acts as closed-loop feedback to the user. **b**. Example reconstructions of cursor position (top) and velocity (bottom) from encoder estimations (dashed black), compared to the actual position and velocity (dashed blue) with a time-shuffled baseline (solid gray). Distributions of average *R*^2^ values between actual and reconstructed position (top right) and velocity (bottom right) for actual data versus time-shuffled controls (N = 14, median; shading shows the 25th - 75th percentile; Two-sided Wilcoxon signed-rank test, (position) **p= 0.0001, (velocity) **p= 0.0001). **c**. Product of the decoder matrix with feedforward contributions of the encoder matrix (N = 14, median; shading shows the 25th - 75th percentile). Black dashed lines represent values that yield perfect trajectory tracking. **d**. Product of the decoder matrix with feedback contributions of the encoder matrix (N = 14, median; shading shows the 25th - 75th percentile). Black dashed lines represent values that yield closed-loop stability. **e**. Average change in angle between final and initial encoders (∠(ℛ (*E*_*f*_ ), ℛ (*E*_*i*_))) (dark purple) and consecutive encoders (∠(ℛ (*E*_*t*+1_), ℛ (*E*_*t*_))) (light purple). Average change in angle between final and initial decoders (∠(𝒩 (*D*_*f*_ )^⊥^, 𝒩 (*D*_*i*_)^⊥^)) (dark orange) and consecutive decoders (∠(𝒩 (*D*_*t*+1_)^⊥^, *𝒩* (*D*_*t*_)^⊥^)) (light orange) (N = 14, center shows median; box shows 25th–75th percentiles; whiskers extend to 1.5 × this interquartile range; two-sided Wilcoxon signed-rank test; (left)**p= 0.00012, (right)**p= 0.00012) **f**. Average change in angle between final and initial encoders (∠(ℛ (*E*_*f*_ ), ℛ (*E*_*i*_))) (dark purple) and final and initial decoders (∠(𝒩 (*D*_*f*_ )^⊥^, *𝒩* (*D*_*i*_)^⊥^)) (dark orange) as a function of trial number (N = 14, median; shading shows the 25th - 75th percentile). **g**. Average change in angle between consecutive encoders (∠(ℛ(*E*_*t*+1_), *ℛ* (*E*_*t*_))) (light purple) and consecutive decoders (∠(𝒩 (*D*_*t*+1_)^⊥^, *𝒩* (*D*_*t*_)^⊥^)) (light orange) over the course of a trial (N = 14, median; shading shows the 25th - 75th percentile).

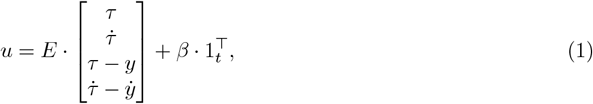

where 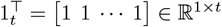, and *t* indicates time.

The encoder matrix has both feedforward (*F* ) and feedback (*B*) components:

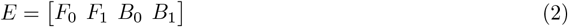

such that each component of task information, 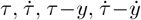, respectively corresponds with an element of the encoder, *F*_0_ ∈ℝ^64×2^, *F*_1_ ∈ ℝ^64×2^, *B*_0_ ∈ℝ^64×2^, *B*_1_ ∈ ℝ^64×2^. The subscripts 0 and 1 represent the 0^*th*^- and 1^*st*^-order dynamics of position and velocity, respectively. The output of the encoder is an EMG activity time series *u* ∈ℝ^64×*t*^ which is input to the decoder, *D* ∈ ℝ^2×64^. The decoder outputs a cursor velocity, which is integrated by the system dynamics *M* to a cursor position *y*. We estimated the user’s encoders *E* ∈ ℝ^64×8^ within a trial with linear regression using 20 second batches of data, corresponding to the time intervals where the decoder was held constant (see Methods). This model appeared to capture useful aspects of user behavior, yielding reconstructed cursor position and velocity trajectories that were correlated with the actual cursor movements (Fig. 2B).

We then used our encoder model to quantify decoder-encoder interactions and analyze performance of the closed-loop system. One common challenge in co-adaptive systems is convergence to a stable solution or performance [25, 27, 28]. Our performance-based analysis above suggested that our co-adaptive interface may have converged (Fig. 1C,D), but does not prove stability nor ideal performance. We exploited the separation of the feedforward and feedback pathways in our encoder model to empirically measure the stability and tracking error properties of the closed-loop encoder-decoder system. Perfect trajectory tracking is obtained when the user’s feedforward input *F* becomes the pseudo-inverse of the decoder *D* and system dynamics *M*, which results in the following conditions: *D* · *F*_0_ = 0 and *D* · *F*_1_ = *I*, where 0 is the zero matrix and *I* is the identity matrix (see Methods, Equation (13)). We used the known decoder values and estimated user encoders to quantify these values in our experimental data, which revealed that *D* · *F*_1_ approach the analytically-ideal values on average across all trials (Fig. 2C). Interestingly, we found that *D* · *F*_0_ slighly deviated from the expected values for perfect tracking (Fig. 2C). These deviations from analytical ideal were found to correlate with user tracking error (Extended Data Fig. 4), suggesting that our encoder estimates capture meaningful properties of user behavior. Similarly, closed-loop stability can be achieved if the feedback components of the decoder-encoder product (*D* · *B*_0_ and *D* · *B*_1_) are approximately negative-definite diagonal matrices. In this case the eigenvalues of the closed-loop system’s dynamics are approximately located at the diagonal entries (i.e. in the left half complex plane), ensuring stability [50, Chapter 5]. Our experimental data observations matched the predictions for a stable system (Fig. 2D).

We also used our encoder estimation model to revisit user-learning-related changes in experiments. We found that user encoder subspaces showed larger changes across the 5-minute trial compared to subsequent time-intervals (Wilcoxon signed-rank test, *p <* 0.001) (Fig. 2E, left; see Methods), consistent with directed changes observed in the EMG tuning curves (Fig. 1H, I). As expected, decoder subspaces also showed directed changes within each trial (Wilcoxon signed-rank test, *p <* 0.001) (Fig. 2E, right; see Methods). We further observed that the within-trial change in user encoder subspaces decreased across sequential trials (Fig. 2F), mirroring our EMG and performance-based analyses showing that users also learn over the course of the experiment (Fig. 1J, K). Decoder changes within each trial, in contrast, did not notably vary across trials in the experiment since the decoders were re-initialized randomly at the beginning of each trial (Fig. 2F). Our models therefore revealed evidence of co-adaptation between the encoder and decoder that was consistent with our EMG-based analyses.

Finally, encoder and decoder changes within each trial, paired with system-level analyses, revealed nuances of how the co-adaptive system converged (Fig. 2G). The amount of change between consecutive decoders starkly decreased over the course of a trial, consistent with the algorithm converging towards a solution from its random initialization. User encoders, in contrast, changed a similar amount over the trial. This is consistent with users learning across trials, where they do not “re-initialize” their encoder on each trial. Importantly, both encoders and decoders do not converge towards zero change within the five minute trial, despite the fact that encoder-decoder pairs (Fig. 2C,D) and task performance (Fig. 1K) appear to converge. Considered together, our analyses suggest that the joint user-decoder system converges, yet neither agent becomes completely stationary. By modeling the user, our control-theoretic encoder model allowed us to more precisely resolve user-decoder interactions within the closed-loop, co-adaptive interface.

### 2.3 Game theory model generated predictions for co-adaptation outcomes

Having established an experimental platform for co-adaptive interfaces and validated tools to analyze user-decoder adaptation, we next aimed to build tools to predict how users and decoders will interact. We leveraged game theory to model the co-adaptive interactions that arise between these two agents optimizing their individual objectives [51, 52] (Fig. 3A). The two agents in our experiment—the user and decoder—co-adapt to achieve a shared goal: reduce task error. However, the user likely trades off error with effort, preferring strategies that achieve reasonable performance while reducing muscle activity [53]. The decoder’s adaptation algorithm was analogously designed to trade off error with the magnitude of the transformation (or gain) it defines between EMG inputs and cursor velocity outputs. We therefore defined cost functions for the user’s encoder *E* and the decoder *D* as a linear combination of task error *e* and agent effort *f*,

**Fig. 3.**
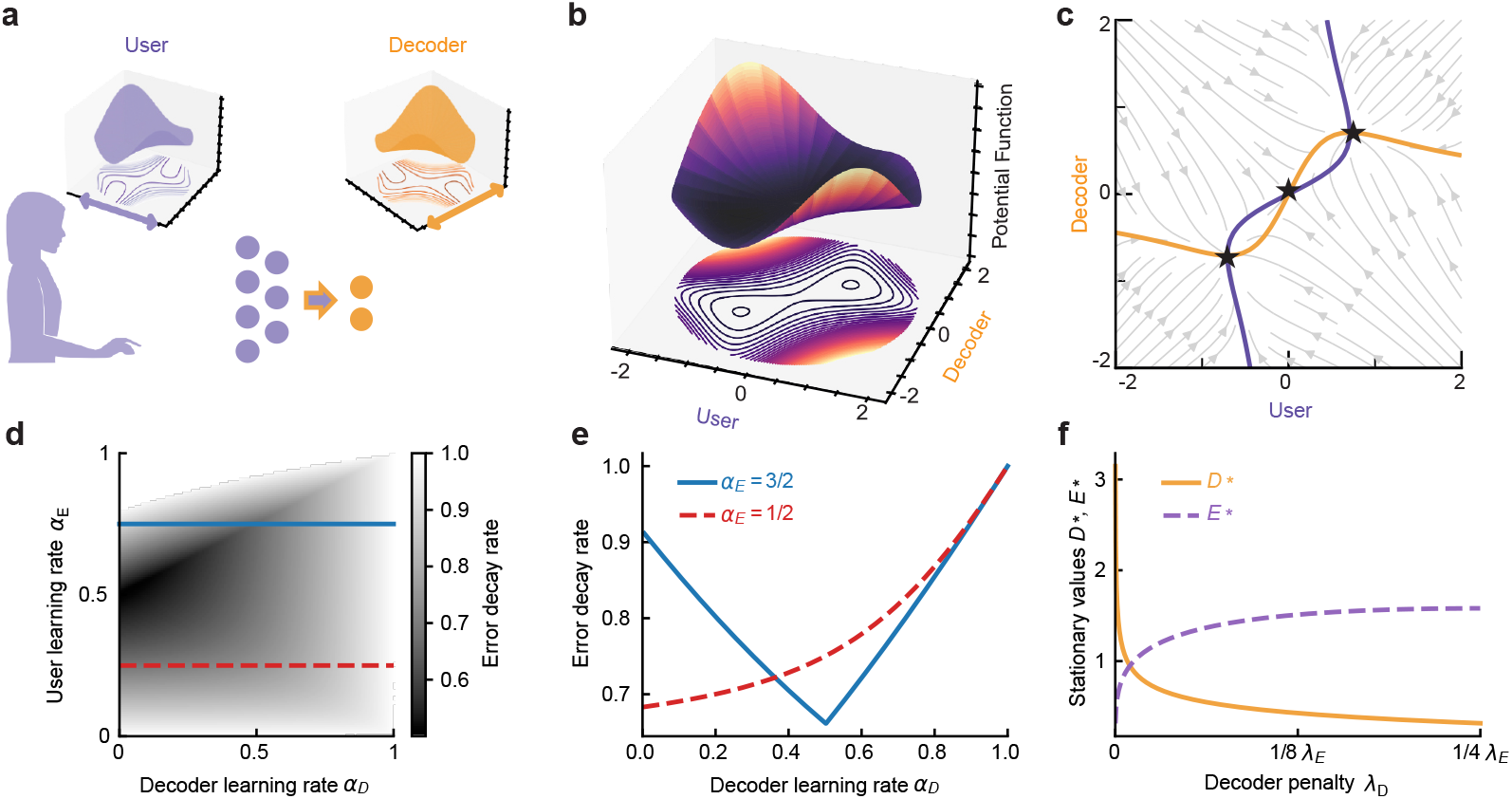
Game-theoretic Model of Co-adaptive User-Machine Systems. **a**. Schematic of co-adaptive interfaces where the user and decoder are modeled as adapting to minimize their own individual cost functions (inset plots for each agent). **b**. Visualization of the potential function that describes dynamics within the user-decoder game model (simplified 1D user and decoder, see Methods). Two axes represent the scalar values of the user and decoder actions, and the vertical axis represents the value of the potential function. A 2D projection of the potential function is also shown. **c**. Gradient field of the user and decoder cost functions (with equal penalty terms, *λ*_*D*_ = *λ*_*E*_ ). Purple (user) and orange (decoder) curves are nullclines (where the agent’s gradient equals 0), which intersect at stationary points (black stars). **d**. Heat map showing error decay rate as a function of learning rates in Equation (5) with penalty parameters *λ*_*D*_, *λ*_*E*_ = 1*/*4. **e**. Error decay rate as a function of decoder learning rate (*α*_*D*_) for two values of encoder learning rate (*α*_*E*_ = 3*/*2, blue, and *α*_*E*_ = 1*/*2, red) corresponding to horizontal slices of panel D. **f**. Stationary values of scalar decoder (orange) and encoder (purple) as a function of decoder penalty parameter (*λ*_*D*_) relative to the value of encoder penalty (*λ*_*E*_ ).

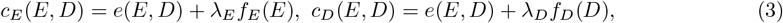

where *λ*_*E*_ and *λ*_*D*_ are effort penalties. For tractability, we limit our analysis in this section to the 1-dimensional case where *E* and *D* are scalars as in [54]. In our simplified case, the error in our task is the square difference between the feedforward decoder output and target velocities, 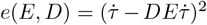. We similarly defined effort as *f*_*E*_(*E*) = *E*^2^, *f*_*D*_(*D*) = *D*^2^.

The structure of these cost functions define a potential game [55] (Fig. 3B), which can be analyzed in terms of a single potential function,

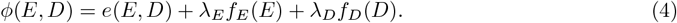

Using this potential function, we can characterize the existence of and convergence to stationary points (*E*_*_, *D*_*_) defined by the conditions *∂*_*E*_*ϕ*(*E*_*_, *D*_*_) = 0 and *∂*_*D*_*ϕ*(*E*_*_, *D*_*_) = 0. These game-theoretic stationary points represent joint strategies in the co-adaptive system that persist for a prolonged period of time. Such stationary points in a neural interface could correspond to stable neural representations (neural “encoding”) and stable decoder parameters, which have been observed in co-adaptive BCIs [2, 10, 14, 27, 40].

We model encoder and decoder adaptation in discrete time using gradient descent for *E* and smoothed best response for *D*,

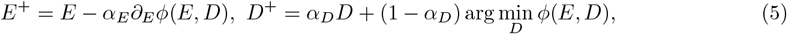

where *α*_*E*_ *>* 0, *α*_*D*_ ∈ (0, 1) are learning rate parameters. The update for *D* is consistent with what is implemented in the experiments (see Methods) and was based on the methods in [10, 40]. We propose a gradient-based update for *E* as a model for user learning that incrementally improves performance, which is consistent with our findings for how encoders change (Fig. 2E-G).

Analyzing game dynamics (see Methods for calculation details) shows that the existence of stationary points is unaffected by learning rates *α*_*E*_, *α*_*D*_, but these parameters determine the system’s overall rate of convergence—or, indeed, whether convergence occurs at all (Fig. 3D, Fig. 4A). When *λ*_*E*_ and *λ*_*D*_ are less than unity, there are three stationary points (*E*_*_, *D*_*_): one at the origin and a mirror pair (see Methods for analytical expressions). The stationary point at the origin is a saddle, and the other two are local minima of the agents’ shared potential function (see Methods). Existence of multiple stationary points in the co-adaptive model suggests the same for the experimental co-adaptive system, and initialization of either agent could affect the stationary point the system converges to. This prediction is tested in Extended Data Fig. 5.

**Fig. 4.**
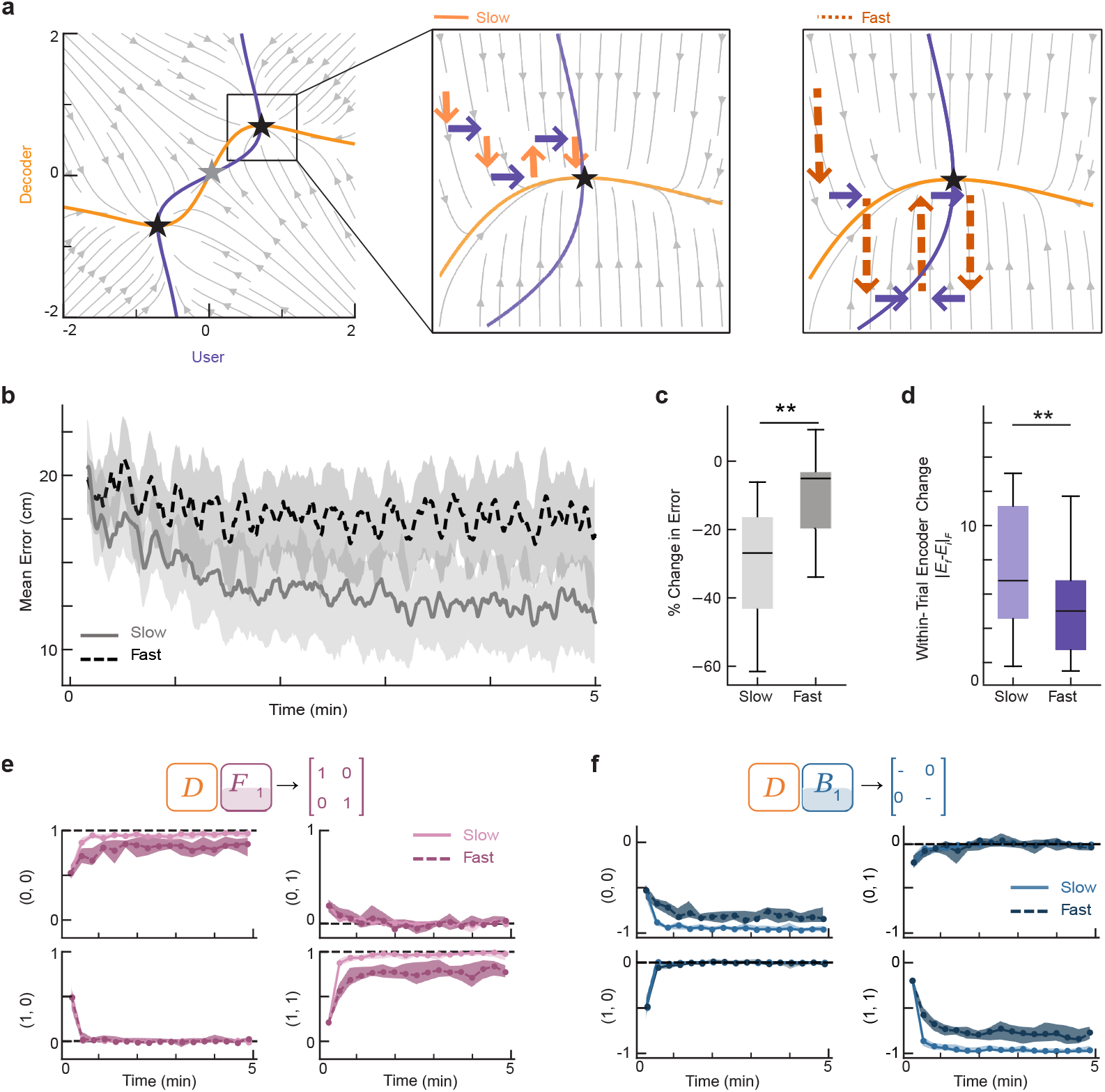
Decoder learning rate influences interface performance and user learning. **a**. Model predictions. All panels show the gradient field of the user and decoder cost functions. Purple (user) and orange (decoder) curves show nullclines (where the agent’s cost gradient equals 0) that intersect at stationary points (black stars). Slow (middle) and fast (right) panels zoom in on one stationary point and illustrate changes in the decoder (shades of orange) and encoder (purple) for different decoder learning rates. **b**. Average error as a function of time within the 5-minute trial for fast (dashed dark orange) and slow (solid light orange) decoder learning rates (N = 14, median; shading shows the 25th - 75th percentile). **c**. Percent change in error from the start of the trial (first 30 seconds) to the end of the trial (last 30 seconds), separated by fast (dashed black) and slow (solid gray) learning rates (N = 14, center shows median; box shows 25th–75^th^ percentiles; whiskers extend to 1.5 × this interquartile range; two-sided Wilcoxon signed-rank test, **p = 6.1*e* − 5). **d**. Average change in user’s encoder within trials (∥*E*_*f*_ −*E*_*i*_ ∥) for the slow and fast decoder learning rate conditions (N = 14, center shows median; box shows 25th–75th percentiles; whiskers extend to 1.5 × this interquartile range; two-sided Wilcoxon signed-rank test, **p= 1.8*e* − 4). **e**. Product of the decoder matrix with first-order feedforward (*F*_1_) contributions of the encoder matrix (N=14, median; shading shows the 25th - 75th percentile) for fast (dashed dark pink) and slow (solid light pink) decoder learning rates. Black dashed lines represent values that yield perfect trajectory tracking. **f**. Product of the decoder matrix with first-order feedback (*B*_1_) contributions of the encoder matrix (N=14, median; shading shows the 25th - 75th percentile) for fast (dashed dark blue) and slow (solid light blue) decoder learning rates. Black dashed lines represent values that yield closed-loop stability.

**Fig. 5.**
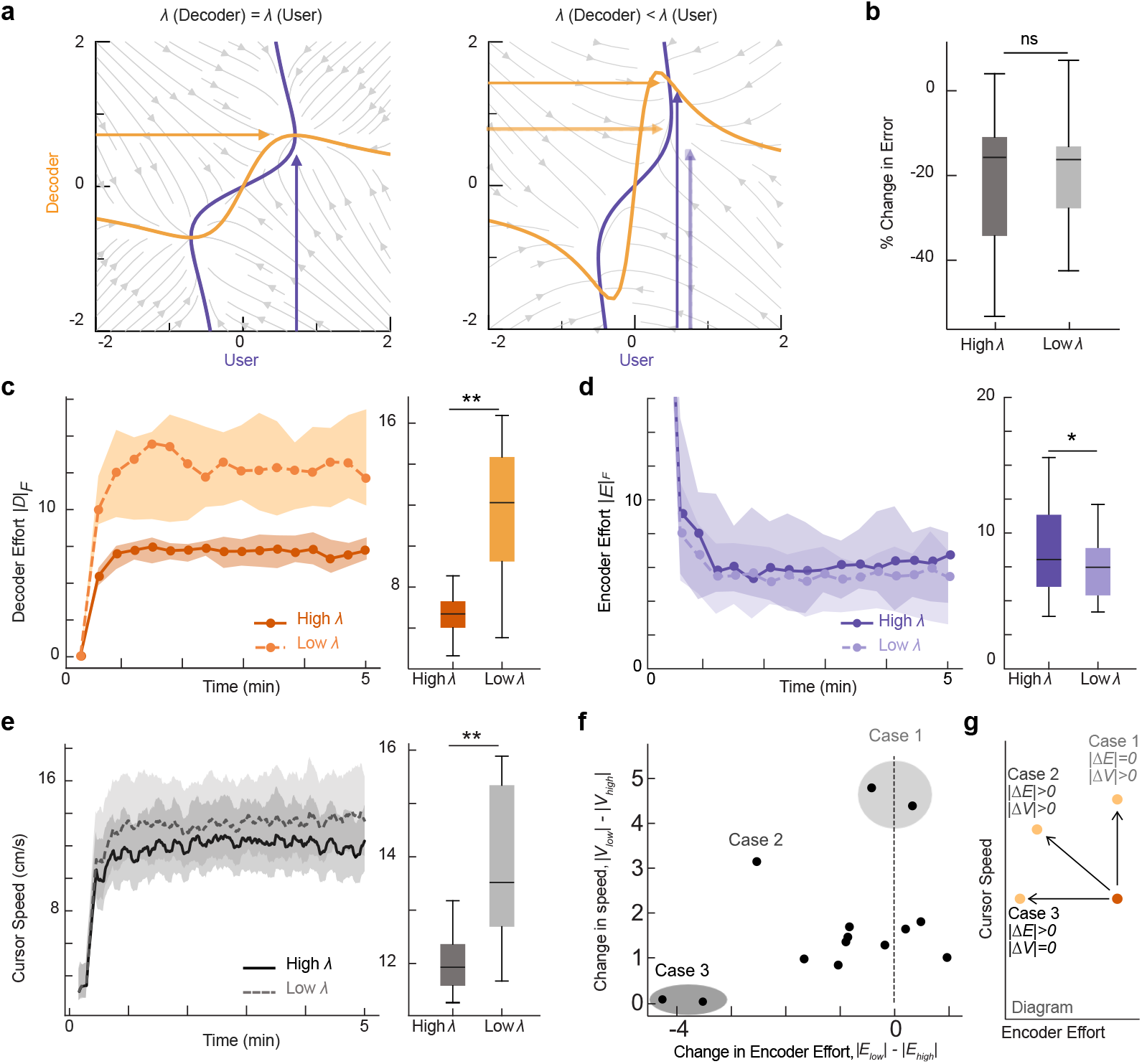
Decoder penalty affects user effort and cursor velocity but not performance. **a**. Model predictions. Panels show gradient fields of the user and decoder cost functions (format as in Fig. 4A) when the decoder penalty term is equal to (left) or less than (right). Arrows depict expected stationary points of user (purple) and decoder (orange). **b**. Percent change in error from the trial start (first 30 seconds) to end (last 30 seconds), separated by high (dark gray) and low (light gray) decoder penalty term conditions (N=14, median; shading shows the 25th - 75th percentile; two-sided Wilcoxon signed-rank test, ns = 0.17). **c**. Left: Average magnitude of the decoder matrix (norm) over time in the trial for low (dashed light orange) and high (solid dark orange) decoder penalty terms (N = 14, median; shading shows the 25th - 75th percentile). Right: Box plots are average decoder effort across the trial for each subject (N=14, center shows median; box shows 25th–75th percentiles; whiskers extend to 1.5 × this interquartile range; one-sided Wilcoxon signed-rank test, **p = 6.1*e* − 5). **d**. Left: Average magnitude of the user encoder matrix (norm) over time in the trial for low (dashed light purple) and high (solid dark purple) decoder penalty terms (N = 14, median; shading shows the 25th - 75th percentile). Right: Box plots are average effort for each subject across the trial (N = 14, center shows median; box shows 25th–75th percentiles; whiskers extend to 1.5 × this interquartile range; one-sided Wilcoxon signed-rank test, *p= .018). **e**. Left: Average cursor speed as a function of time in the trial for low (dashed light gray) and high (solid dark gray) decoder penalty terms (N = 14, median; shading shows the 25th - 75th percentile). Right: Boxplots are average cursor speed across the trial for each subject (N=14, center shows median; box shows 25th–75th percentiles; whiskers extend to 1.5 × this interquartile range; one-sided Wilcoxon signed-rank test, **p = 6.1*e* − 5). **f**. Average difference in cursor speed between the low and high decoder penalty conditions plotted against the average difference in encoder efforts between the low and high decoder penalty conditions for each individual participant (black dots). Shading to illustrate qualitatively identified groupings. **g**. Schematic illustrating the relationship between encoder effort and cursor speed when decoder effort changes. Dots represent encoder effort and velocity for high (dark orange) and low (light orange) decoder penalty. Arrows depict the potential scenarios of how decoder penalty can shift encoder effort and velocity.

Setting learning rates of adaptive decoders has attracted significant interest in part because of its notable impact on overall neural interface performance [7, 25, 27, 28, 40]. Our model allows us to make principled predictions that could be used to design and optimize adaptive algorithms. We assessed convergence of Equation (5) to the stationary points (*E*_*_, *D*_*_) with *E*_*_, *D*_*_≠0 by linearizing the dynamics about that point and then computing the eigenvalues of this linearization [56, Chapter 5]; the linearization was the same for both stationary points. We refer to the larger eigenvalue as the *error decay rate*, as it is the smallest number *ρ >* 0 such that the error *e*(*i*) is bounded above by *e*(0)*ρ*^*i*^ where *i* is iteration number. Interestingly, this analysis predicts a “goldilocks” effect of decoder learning rate on system convergence, consistent with past models [25], for one regime of user learning; but predicts a monotonic relationship between system convergence and decoder learning rate for other user learning rate regimes (Fig. 3D,E). This complex landscape illustrates that empirical optimization is challenging and demonstrates the value of our computational framework for theory-guided optimization. The prediction that learning rate affects error decay rate is tested in Fig. 4.

We then explored the contribution of penalty terms *λ*_*E*_, *λ*_*D*_, which influence user-decoder phase portraits (Fig. 5A). Altering these penalties shifts the stationary points (*E*_*_, *D*_*_) and, thus, affects the user and decoder efforts, *f*_*E*_ and *f*_*D*_. We find that an inverse relationship holds between *E*_*_ and *D*_*_ for a range of *λ*_*E*_ and *λ*_*D*_: fixing *λ*_*E*_ and varying *λ*_*D*_ from 0 to 1*/*4*λ*_*E*_, *D*_*_ decreases monotonically and *E*_*_ increases monotonically (Fig. 3F). This prediction is tested in Fig. 5.

To summarize, our game theoretic model generates the following predictions about user-decoder co-adaptation:

1. Overall system convergence and performance are influenced by both user and decoder learning rates.
2. Decoder penalty terms affect the location of stationary points in the overall system, which determines user effort. This finding predicts that users and decoders, effectively, “trade-off” effort to maintain system performance.
3. Because multiple stationary points exist, each learner’s initialization will affect the final stationary point reached by the system. This predicts that decoder initialization will influence a user’s final learned encoder.

### 2.4 Decoder learning rates can disrupt co-adaptation

Our game-theoretic model emphasizes the importance of both the user and decoder learning rates for convergence to stationary points. One agent adapting faster than the other may limit or prevent convergence to stationarity (Fig. 4A). Our model predictions echo results from previous models [25] and experimental findings from both invasive neural and non-invasive body-machine interfaces [7, 57]. We examined the validity of our model’s predictions by testing the effect of decoder learning rates in our co-adaptive myoelectric interface system. Because the user’s learning rate was unknown and could not be estimated *a priori*, we varied the decoder learning rate, testing two extremes: slow and fast (see Methods). Our model makes two alternate predictions for the outcome of this manipulation, depending on user learning rate: if users learn relatively rapidly, these two extremes should yield similar performance; if users learn more slowly, fast decoder learning rates should reduce performance (Fig. 3E). Decoders for both learning rate conditions were initialized randomly, matching all other conditions, and then adapted according to the specified cost function. Task performance was worse for co-adaptive interfaces where the decoder adapted with the fast learning rate compared to the slow learning rate throughout all trials (Fig. 4B), and performance improved more within the trial in the slow condition compared to the fast condition (Wilcoxon signed-rank test, *p <* 0.001) (Fig. 4C).

We found that the decoder learning rate also affected the user’s encoders. We used our encoder estimates (Fig. 2) to quantify how much users’ encoders changed over the course of the 5-minute trial in each decoder condition (see Methods). User encoders changed less within trials when the decoder adapted fast compared to slow (Wilcoxon signed-rank test, *p <* 0.001) (Fig. 4D), suggesting that the decoder may have adapted too fast for the user’s encoder to match. We then analyzed user-decoder interactions, which confirmed that rapid decoder adaptation disrupted co-adaptation. The decoder-encoder pairs in the slow condition approached the system ideal for perfect trajectory tracking and closed-loop stability, but the fast condition decoder-encoder pairs deviated from the system ideal state in both the first-order feedforward (Fig. 4E) and feedback conditions (Fig. 4F), showing that the fast learning rate negatively impacted the co-adaptive interactions for both trajectory tracking and closed-loop stability. Our experimental results corroborated our game-theoretic prediction that decoder learning rates will influence system behavior, though precise validation experiments will require improved methods to estimate user learning rates (see Discussion).

### 2.5 Decoder “effort” penalty influenced user “effort” without changing performanc

Our game-theoretic framework includes the regularization terms, ∥*D*∥ and ∥*E* ∥, which correspond to what we refer to as “effort”. Including regularization terms enforces convergence to a stable stationary point (See Game Theory Methods). In the model, the scaling of the decoder regularization term, controlled by *λ*_*D*_, influences the decoder effort. In turn, the user effort is also influenced by *λ*_*D*_ due to the linear relationship between the decoder and cursor velocity (Eq. 1). As the decoder penalty changes in our model, the stationary point and encoder effort correspondingly shift (Fig. 5A). Our model therefore predicts that changing decoder effort will not impact convergence to stationarity in the co-adaptive interface, but will influence user behavior by shifting their effort. Past work suggests that users can adapt their encoder to “trade-off” with a decoder in a co-adaptive interface where the decoder only prioritized optimizing task performance [10]. Yet, many studies highlight limitations on the degree of user flexibility in a range of neural interfaces [58, 59]. It is therefore unknown whether users will flexibly trade-off with a decoder that aims to optimize multiple objectives, including its own individual “effort”.

We tested the impact of two penalty terms – low penalty and high penalty – on decoder-encoder co-adaptation in our myoelectric interface experiments. Decoders for both penalty conditions were initialized randomly, matching all other conditions, and then adapted according to the specified cost function (see Methods). Performance was not impacted by the penalty terms (Wilcoxon signed-rank test, *p >* 0.05) (Fig. 5B), suggesting the co-adaptive systems converged similarly for both decoder conditions. Decoder effort was influenced by the penalty term as expected: higher decoder penalty resulted in decreased decoder effort (Wilcoxon signed-rank test, *p <* 0.001) (Fig. 5C). As predicted by our model, the decoder effort affected the encoder effort to reflect the trade-off in effort between the two learners. Thus, higher decoder penalty terms resulted in increases in the encoder effort (quantified by the norm of the *F*_1_ component of the encoder; Wilcoxon signed-rank test, *p <* 0.05)(Fig. 5D). These results held regardless of the learning rate of the decoders (Extended Data Fig. 6).

While our experimental results broadly matched model predictions, we noticed that the relative decrease in encoder magnitude with respect to the increase in decoder magnitude was smaller than predicted by the theory (Fig. 5C vs. 5D). Importantly, decoder “effort” controls the magnitude of *D*, which defines the gain between user EMG and cursor velocity. If users do not change their effort to perfectly mirror changes in the decoder effort, the cursor’s speed - an aspect of task performance not captured in our model and not directly linked to task performance - would change. We found that decoder penalty terms influenced the cursor speed during tracking, with a lower decoder penalty term leading to faster cursor speeds on average across users (Wilcoxon signed-rank test, *p <* 0.001)(Fig. 5E). Interestingly, these trade-offs differed across our participants (Fig. 5F), potentially capturing individual preferences. We noticed clear outlier participants who chose to decrease their effort and maintain a similar cursor speed, or alternately maintain a similar effort to go faster; the remainder of our participants changed both effort and speed to varying degrees. This suggests three broad groups of strategies adopted by users (Fig. 5G). These different strategies, however, did not correlate with task performance, suggesting that users may identify compromise solutions to trade-off with the decoder without sacrificing task performance. These findings highlight experimental deviations from our model predictions for strict encoder-decoder tradeoffs. We also found only subtle evidence to support our model’s predictions related to the influence of decoder initialization (Prediction 3; Extended data Fig.5; see Discussion). Despite these deviations, our experimental results broadly confirmed our game-theoretic model predictions and highlighted the potential of decoder adaptation to influence the user’s encoder during closed-loop co-adaptation.

## 3 Discussion

We extended quantitative modeling techniques from control theory and game theory to reveal how decoding algorithms can shape user adaptation in neural interfaces. The critical first step was developing a myoelectric interface testbed that enabled systematic experiments examining how users learn to control an unfamiliar interface alongside different adaptive algorithms. Our platform provides advantages over existing human-in-the-loop simulators and related approaches, which may not fully capture how people respond to adaptive algorithms in closed-loop settings [60–64].

We then created control theory-based estimations of user encoder transformations that allowed us to quantify critical closed-loop properties like trajectory-tracking and stability (Fig. 2C,D) and illustrated how decoder design impacted these properties (Fig. 4E,F). Prior work has shown that encoder-decoder stability likely contributes to interface performance and user learning [2, 7, 10], but efforts to quantify stability in adaptive neural interfaces have focused on the decoder alone [27, 28]. By developing methods to analyze encoder-decoder pairs, we were able to characterize stability in the full co-adaptive system (Fig. 2, Fig. 4E,F) and reveal how decoding algorithms can influence user adaptation. While our approach used a simplified model of user closed-loop control policies, our results highlight the promise of these frameworks for designing and analyzing neural interface systems. Expanding the accuracy of our models, by incorporating details like sensory processing delays, will likely lead to further advances.

We also developed a game-theoretic model to generate predictions about the behavior of different co-adaptive systems. Designing co-adaptive interfaces requires confronting a vast space of algorithm parameters that can influence decoder and interface performance [7, 25, 27, 40, 65]. Our approach provides experimentally-validated methods to quantitatively model and predict outcomes of co-adaptive dynamics. These tools will enable principled interface design and optimization by considering multiple objectives and user-decoder interactions. Current neural interfaces largely aim to maximize performance on a task without considering other aspects of interface performance, such as stability or robustness. Our results demonstrate an example of training algorithms to consider multiple objectives, balancing task error with decoder effort, which in turn influences user effort (Fig. 5). Future interfaces might be designed to consider additional objectives like stability, robustness, or performance across multiple tasks. We foresee our game-theoretic optimization-based framework becoming especially useful as the functionalities of neural interface tasks grow.

Our game-theoretic model is particularly valuable for predicting user-decoder interactions that influence system performance, such as decoder learning rates (Fig. 3D-F). We found that a faster decoder learning rate yielded worse overall performance (Fig. 4C) and altered trajectory-tracking and stability in encoder-decoder pairs (Fig. 4E,F) in our experimental myoelectric interface. Our results add to prior experimental observations from co-adaptive kinematic [7, 25] and brain-computer interfaces [25, 57]. Interestingly, many high-performance adaptive algorithms in invasive neural interfaces adapt rapidly [13, 28, 66]. Other co-adaptive interface models and experiments suggest intermediate learning rates are optimal [25]. We found that fast decoder learning rate condition disrupted co-adaptive dynamics in our interface, and our data suggest a more monotonic relationship between decoder learning rates and system performance. Our model highlights that the learning rates of both the user and decoder determine system dynamics (Fig. 3D-F) and our experimental data are most consistent with a regime where users learned slowly. While we cannot yet verify user learning rates, it is plausible that our naïve participants may have adapted at different rates than the more experienced participants in invasive brain-computer interface studies and participants using well-learned input interfaces (e.g., a computer mouse) in a coadaptive system [25]. Indeed, user learning rates may vary across different bio-signal modalities [20]. Our framework provides a path to optimize neural interfaces by tailoring the algorithm adaptation rates based on user behavior. This raises important future challenges to develop methods that can perform on-the-fly estimates of critical properties of user behavior, such as learning rate.

Our game theory model and experimental results also revealed user-decoder interactions that influence user behavior, such as decoder “effort” (Fig. 5D). This demonstrates the feasibility of using our framework to optimize interfaces for particular goals. In myoelectric interfaces, manipulating decoders to encourage users to learn encoders that use less muscle activity could be desirable to maximize interface usability for extended periods. Alternately, decoders that encourage users to increase muscle activation towards a desired target could be desirable for motor rehabilitation applications [23, 67]. Interestingly, we saw that users responded differently to varying decoder penalty terms, potentially reflecting personal preferences for effort versus cursor speed (Fig. 5F). Customizing interfaces based on dynamic user preferences has been shown to be beneficial in exoskeleton interfaces [68]. Individual preferences for cursor dynamics have also been seen in invasive neural interfaces [69]. Extending our game-theoretic frameworks to include user preferences will enable design of fully personalized, custom neural interfaces, which may make these technologies more widely accessible [70].

Our game theory model highlighted that multiple decoder-encoder stationary points exist in neural interfaces. This raises the possibility that the initial decoder can affect the final stationary point and, in turn, the final user encoder, without significantly impacting performance (Extended Data Fig. 5A). Our experimental results confirmed that performance was not impacted by decoder initialization (Fig. 1E), consistent with prior work testing random decoder initializations in neural interfaces [40]. While performance was not different, we found that initialization did subtly impact the stationary points as identified by the final decoder-encoder matrices (Extended Data Fig. 5E). Our findings suggested that decoder initialization might bias the encoders learned by users, which could be particularly beneficial for rehabilitation applications, which aim to shape users’ behavior towards a particular goal. Our results also suggest that it may be critical to carefully consider initial training protocols for any neural interface, as it may influence user learning trajectories.

Our experiments revealed occasional deviations from model predictions, including deviations from strict encoder-decoder trade-offs (Fig. 5F) and only subtle impacts of the decoder initialization (Extended Data Fig. 5). This finding suggests that the actual user’s cost might differ from our model’s assumptions. For instance, our model predicts that decoder initialization would lead users to completely invert the relationship between EMG/movement and cursor velocity (Extended Data Fig. 5A), which ignores the likely possibility that users have some bias in strategies to control the interface. Additionally, there may be multiple ways to represent “effort”, and these definitions may also become unclear in less embodied invasive neural interfaces. Prior work has considered that users minimize effort [53, 71] via muscle activation [72], metabolic cost [73, 74], and motor variability [75]. Studies into how users learn to control neural interfaces suggest that other features, such as correlations between neurons or muscles, may be key factors [58, 59]. Deeper insights into how users learn to control different kinds of interfaces will improve our ability to model their cost functions to predict user-decoder interactions.

By developing computational and experimental methods to study co-adaptive systems, our results reveal how interface algorithms can influence user behavior and system performance. Specifically, our model and experimental findings show that decoder learning rate, penalty terms, and initialization influence encoder-decoder dynamics in co-adaptive interfaces. These observations are consistent with recent work suggesting that adaptive algorithms can influence neural representations learned in brain-computer interfaces [76]. Our framework provides paths to harness the complex interactions between users and algorithms in closed-loop interfaces. For example, our results hint at the eventual possibility of designing a “curriculum” to help naive users learn to control neural interfaces, which is a task that some find challenging [47]. Decoder learning rates may need to be significantly slower early in learning when users widely explore control strategies [77–79] and adapt as users master the interface. Strategic decoder manipulations during a user’s initial exploration could be designed to bias users towards particular strategies. We can borrow concepts from training neural networks [80] to envision a decoder adaptation curriculum to intelligently shape user learning, fully harnessing the power of co-adaptive interfaces. Our framework provides a versatile and principled way to model and design the next generation of smart, personalized adaptive neural interfaces.

## Methods

### Experimental Methods

#### Participants

Fourteen volunteers were recruited for this study and gave their written consent prior to experiments, according to study procedures approved by the University of Washington’s Institutional Review Board (IRB #STUDY00014060). All participants were compensated monetarily for their time. All participants had no known motor disorders. Participant demographic information (including gender, weight, height, age, and handedness) was collected via a demographics survey that participants completed before the experiment (see Table 1). Only forearm circumference was measured by the experimenter, all other information was self-reported.

#### Experimental design

Participants were asked to control a cursor on the screen – using muscle activity from their forearms measured via electromyography (EMG) – to follow a 2-D continuous target trajectory as closely as possible (Extended Data Fig. 1A). Participants were told that they might not be able to control the cursor at the beginning of the trial but to expect that their cursor control would improve as the trial progressed. Participants received no other cues or information about the decoder or study conditions. If the participant’s cursor was stuck in the corner or edge of the screen for longer than 3.33 seconds, the cursor was automatically reset to the center of the screen. Participants sat in a chair with no restraints facing a computer screen (HP Compaq L2206tm, 1900 x 1600, 46.5cm x 24.5cm). The computer screen displayed a red target circle (RGB: 1, 0, 0). The cursor was a blue circle (RGB: 0, 0, 1). The target was three times the area of the cursor (Extended Data Fig. 1A).The target and cursor positions were updated and displayed at 60 Hz. The task was programmed using the pygame and LabGraph Python packages.

Following prior work [38, 39, 48], the target traversed a pseudorandom trajectory, which was generated by a sum-of-sinusoids with randomized phases with frequencies that were prime multiples of 0.05 Hz (x-axis frequencies = 0.10 and 0.25 Hz, y-axis frequencies = 0.15 and 0.35 Hz). We randomized the phase of the sum-of-sines so that the reference trajectory would be unpredictable to the participants. Distinct prime multiples were chosen in each direction to provide separability in the x- and y-axes for any frequency-based analysis. To ensure constant signal power, the magnitude of each frequency component was normalized by the frequency squared. The trajectory was different every trial. Each trial was 5 minutes with a 5-second ramp period during which the cursor and target speed slowly increased from stationary to the experimentally-prescribed speeds. This ramp period followed prior experiments [48] and was added to give participants time to recognize the starting cursor and target movements. Participants completed 16 trials of 5-minutes each in 2 blocks; each block consisted of 8 trials. Participants were given a 5-minute break in between each block but their EMG array and placement did not change throughout the experiment.

#### EMG signal collection and preprocessing

EMG signals were obtained using a Quattrocento system (Bioelettronica, Italy). A 64-channel high-density surface EMG electrode array (8mm inter-electrode spacing, 5x13 electrode rectangular layout) was placed on the dominant forearm of each participant, targeting the Extensor Carpi Radialis (Extended Data Fig. 1B). Electrodes were placed on the dominant arm for each participant. Once placed, the electrode array was wrapped with Coban self-adherent wrap (3M, Saint Paul, Minnesota). The electrode cables from the Quattrocento to the array were secured to minimize motion artifacts.

EMG signals were acquired using Biolite Software (Bioelecttronica, Italy) at 2048 Hz on Differential Mode with the built-in low-pass filter of 130 Hz and a high-pass filter of 10 Hz. We filtered and rectified the EMG data following prior preprocessing techniques to compute the EMG linear envelope [81, 82]. A moving average filter was applied to downsample EMG signals from 2048Hz to 60Hz (Extended Data Fig. 1C).

#### Myoelectric Interface Decoder

Real-time myoelectric control was implemented with a velocity-controlled Wiener filter that output a cursor velocity *v*_*t*_ ∈ ℝ^2^,

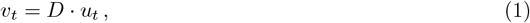

where *u*_*t*_ ∈ ℝ^*N*^ is the processed EMG signals of *N* = 64 channels and *D* ∈ ℝ^2×*N*^ is the decoder mapping. The cursor velocity is integrated to output a cursor position *y*_*t*_ ∈ ℝ^2^,

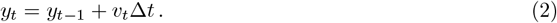

where Δ*t* is the time between cursor updates.

#### Myoelectric Interface Decoder Adaptation

Decoder adaptation consisted of two steps: (1) calculate the optimal decoder *D*^*^ by minimizing the cost function *c*_*D*_ based on the previous 20 seconds of user and trial data:

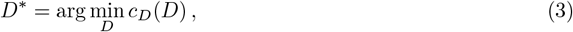

and (2) update the next decoder *D* following the SmoothBatch approach [40], which uses a weighted combination of the prior decoder *D*^−^ and the optimal decoder *D*^*^:

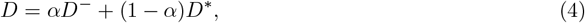

where *α* ∈ (0, 1) is the learning rate. The decoder cost function was minimized using SciPy. To satisfy real-time timing constraints on decoder cost minimization, we initialized 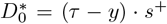 using the previous 20 seconds of data [26]. At the start of each trial, the decoder was initialized by randomizing the decoder weightings from a uniform distribution (using numpy.random.rand) and multiplying by a scalar factor. Each user had two different decoder initializations, referred to as D1 and D2. Initial decoders for each participant’s trial were set to D1 or D2. Note, while we programmed the decoder to update every 20 seconds, due to slight software imprecision, the decoder update actually occurred approximately every 18 seconds.

#### Decoder Cost

The decoder cost function was formulated in earlier work [54] and aims to minimize both the tracking error and decoder effort as the decoder adapts. Minimizing velocity error is a common goal in user-machine interfaces, seen widely across neural interfaces [10, 83], body-machine interfaces [84], and myoelectric interfaces [43, 65]. Decoder effort is considered as part of the decoder cost since our prior theoretical analysis suggested that a regularization term in the user and decoder costs is necessary to ensure convergence in the user-decoder co-adaptation game to stable stationary points [54].

The decoder cost *c*_*D*_ is constructed as a linear combination of the task *error* and decoder *effort* :

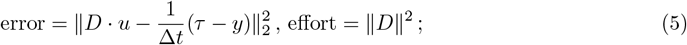

here ∥*x*∥_2_ denotes the 2-norm of signal *x* : [0, *t*] → ℝ^*d*^, and ∥*X*∥ denotes the Frobenius norm of matrix *X* ∈ ℝ^*m*×*n*^. The decoder cost *c*_*D*_ is then,

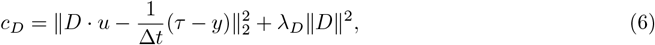

where *λ*_*D*_ is the penalty term of the decoder effort.

#### Decoder Conditions

We varied the decoder cost function and parameters of decoder adaptation to determine how it influenced system performance and user behavior. Specifically, we varied: (1) learning rate *α*, (2) decoder effort penalty term *λ*_*D*_, and (3) decoder initialization. We tested two learning rates: *slow* (*α* = 0.75) and *fast* (*α* = 0.25), two penalty terms: *low* (*λ* = 10^2^) and *high* (*λ* = 10^3^), and two randomized initializations of the *D* decoder matrix (D1 and D2): *positive* (matrix elements chosen uniformly at random in the range [0, 10^−2^]) and *negative* (elements chosen uniformly at random in [−10^−2^, 0]). We tested all combinations of learning rates, penalty terms, and initializations, leading to a total of eight different decoder conditions.

### Data Analysis Methods

#### Performance Metrics

Our primary metric for quantifying performance was the tracking error, calculated as the Euclidean distance between the target and the cursor, | *τ* −*y*|_2_. We assessed changes in task performance within a trial by comparing the mean tracking error in the first (early) and last (late) 30 seconds of the trial (excluding ramp-up time). Because participants’ proficiency in tracking varied, we quantified improvements over time by calculating the relative error: 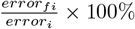, where error_*fi*_ = error_*final*_ - error_*initial*_.

#### Statistical Analyses

All analyses treated participants as individual data points and computed the mean across decoder conditions (learning rate, penalty terms, initialization) and trials. To assess statistical significance across time and conditions for each participant, we used a one- or two-sided Wilcoxon signed-rank test (`scipy.stats.wilcoxon`), which is a paired, non-parametric test. We chose a non-parametric statistical test because of subject-to-subject variability. Figure boxplots were plotted with `matplotlib.pyplot.boxplot` (center line, median; box limits, upper and lower quartiles; whiskers, 1.5x interquartile range; fliers not plotted).

#### EMG Analysis

To quantify the relationship between user EMG activity and cursor movement, we computed EMG direction tuning curves, a well-established analysis used for neural and myoelectric interfaces [42–44].

We first down-sampled and grouped intended user cursor velocities into 10 equally-spaced directions and then fit raw EMG activity to the binned directions to form EMG tuning curves. These EMG tuning curves were created for each participant, each trial, and each EMG signal (Fig. 1F). Note, since the EMG channels are differentially-recorded, we selected 63 EMG signals for the EMG tuning analysis.

To quantify the within-trial EMG changes from early to late trial segments (Fig. 1G) we computed the norm difference between the early and late EMG tuning curves:

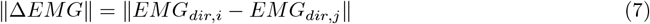

where *EMG*_*dir*_ is the average EMG activity per direction and *i* and *j* indicate time points. All tuning curves were calculated by reconstructing 30-second segments of trial data that equally sampled all cursor directions. To measure how the EMG activity changed across the entire experiment per participant (Fig. 1H), we averaged the |Δ*EMG*|_2_ across all channels for each participant to compute a mean |Δ*EMG*|_2_ per participant per trial.

To find the preferred direction of the EMG activity, we performed cosine-fitting analysis [42]. Specifically, average EMG activity for each intended cursor direction was modeled as:

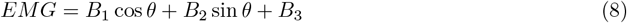

where *θ* represents intended cursor direction and *B*_1_, *B*_2_ and *B*_3_ are model coefficients. The preferred direction (PD) is:

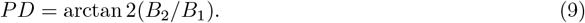

The change between early and late preferred direction (Fig. 1I) was calculated as |Δ*PD*| = |*PD*_*late*_ −*PD*^*early*^ |.

### Encoder Estimation Methods

To model the user’s transformation from task space to user signals, we estimated the user’s encoder *E*. We model the user’s signal as being produced by a combination of feedforward (*F* ) control on position (*τ* ) and velocity 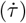 data, as well as feedback (*B*) control on position error and velocity error data ((*τ* − *y*) and 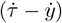, respectively):

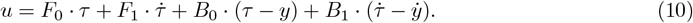

Or, in matrix formulation:

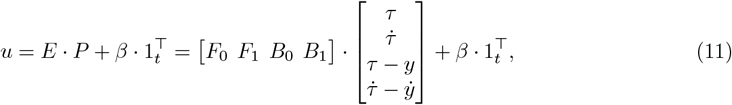

where 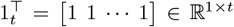, *u* ∈ℝ^*N* ×*t*^, *E* ∈ℝ^*N* ×*d*^, *P* ∈ℝ^*d*×*t*^, *N* is the number of signals, *d* is the dimension of task information, and *β* is an offset factor for estimation (*β* ∈ ℝ^*N* ×1^). In this paper, *N* = 64 EMG channel inputs, *d* = 8 (target position, target velocity, position error, and velocity error for both *x* and *y* dimensions). The selection of *t* determines the timescale on which *E* is estimated.

We numerically estimated users encoders *E* using linear regression of experimentally measured data for *u* and *P* . We estimated *E* every 20 seconds. This corresponds to the timescale on which decoders were updated, thereby generating matched encoder-decoder estimate pairs to analyze co-adaptive dynamics (e.g., Fig. 2C,D). Linear regression was done by constructing a *u* matrix of EMG signals for *t* = 1200 samples (20 seconds of data at 60 Hz) for the 64 EMG electrodes, and a corresponding matrix *P* of position, velocity, position error, and velocity error (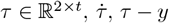. and 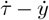, respectively). These data were used to fit equation (11) using linear regression (`sklearn.linearmodel.LinearRegression`) with an intercept offset.

We estimated the accuracy of the myoelectric encoder model by reconstructing EMG signals from the encoder estimation and compared the coefficient of determination (*R*^2^) for the cursor velocity and position decoded from the reconstructed EMG to the actual cursor velocity and position recorded in the trial. To establish a baseline for the predictive power of our encoder model, we also computed the accuracy of cursor velocity and position decoded from time-shuffled EMG data (Fig. 2B). The time-shuffled EMG data was created by permuting the EMG signals within each trial, so that the EMG data came from the same subject and trial but different time points.

### Closed-loop Decoder-Encoder Predictions

We used control theory principles to predict relationships between elements of the encoder *E* and decoder *D*. The decoder is velocity-based, meaning that the cursor position *y* is related to the user’s EMG signal *u* via 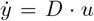 where *D* is the decoder matrix. We model the user’s signal as being produced by a combination of feedforward and feedback control on position and velocity data:

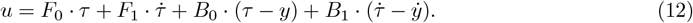

This controller can achieve exact tracking, defined as *y* = *τ*, if *D* · *F*_0_ = 0 and *D* · *F*_1_ = *I* where 0 is the 2 × 2 zero matrix and *I* is the 2 × 2 identity matrix. To see this, suppose at time *t* that *τ* (*t*) = *y*(*t*). Then

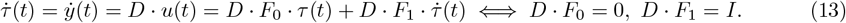

Note that *M* is a simple integrator, so the condition for 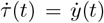 in Equation (13) together with *τ* (*t*) = *y*(*t*) ensures *τ* = *y* at all times. To ensure closed-loop stability, it suffices that *D* · *B*_0_ and *D* · *B*_1_ are negative-definite diagonal matrices [50, Chapter 5].

### Decoder-Encoder Pairs

We calculated decoder-encoder pairs by multiplying the decoder (*D* ∈ ℝ^2×64^) at each update with the feedforward and feedback contributions to the encoder (*E* ∈ ℝ^64×8^) at each update. Each decoder-encoder pair (*D* · *F*_0_, *D* · *F*_1_, *D* · *B*_0_, and *D* · *B*_1_) resulted in a 2×2 matrix representing the dynamics between the encoder and the decoder.

### Encoder Differences

We calculated changes in encoders and decoders over time using the principal angle between subspaces (range space of *E* and orthogonal complement to the null space of *D*) [85] and the norm-difference of encoder matrices: ∥*E*_*i*_ −*E*_*j*_∥, where *i* and *j* indicate time points. Since we started from a decoder that was initialized with randomized weightings and resulted in poor user control, the details of the first encoder *E*_0_ were not readily interpretable. We did not include *E*_0_ in computing or plotting the encoder differences in Fig. 2E - G. To compute encoder changes within a trial, we calculated the difference between the final (*E*_*f*_ ) and initial (*E*_*i*_) encoders, estimated by averaging the first three and last three encoders in a trial, respectively (excluding *E*_0_). To compare data from consecutive encoders to the average change across a full trial (Fig. 2G), we computed the consecutive encoder difference by averaging across three subsequent, non-overlapping encoders and calculating the norm difference. This method ensured that the same number of encoder data points would be compared.

### Game Theory Methods

We modeled the co-adaptive system in a game theory framework involving two agents: the user and decoder. The user and decoder decision variables were the entries in the matrices *E* and *D*, respectively. For tractability, we limit our analysis to the scalar case *E, D*∈ ℝ. Each agent was modeled as having a cost function defined by a linear combination of *task error* and *agent effort*. Agent effort was quantified using the square of the scalars *E* and *D*. Task error was defined as in Equation (5) as the norm squared of *D* · *u* − (*τ* − *y*)*/*Δ*t*. Considering the scalar case and focusing solely on feedback control (i.e. setting *u* = *B*_0_(*τ* − *y*)) yields error (*DB*_0_Δ*t* − 1)^2^(*τ* − *y*)^2^*/*Δ*t*. For simplicity in the analysis we relabel *E* = *B*_0_, normalize by (*τ* − *y*)^2^*/*Δ*t*, and choose coordinates for *D* so that *e*(*D, E*) = (*DE* − 1)^2^ = (1 − *DE*)^2^.

### Stationary points

In the 1-dimensional game formulation, the potential function takes the form

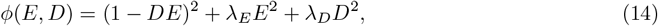

where *D, E* ∈ ℝ. We solved the nonlinear equations *∂*_*E*_*ϕ*(*E, D*) = 0 and *∂*_*D*_*ϕ*(*E, D*) = 0 to compute stationary points as a function of penalty parameters *E*_*_(*λ*_*E*_, *λ*_*D*_), *D*_*_(*λ*_*E*_, *λ*_*D*_), finding a saddle at *E*_*_ = *D*_*_ = 0 and the following mirror pair of local minima:

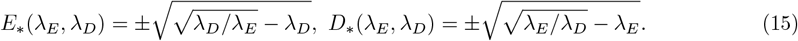

That these points are local minimizers was verified for *λ*_*E*_, *λ*_*D*_ ∈ (0, 1] by numerically determining that the eigenvalues of the Hessian of the potential function *ϕ* were positive at these points.

### Model for adaptation

We modeled agent adaptation using the discrete-time dynamics in Equation (5). To assess convergence of this model for adaptation to stationary points, we linearize the discrete-time dynamics about a stationary point and compute eigenvalues of this linearization. A sufficient condition for stability is that the magnitude of all eigenvalues is less than 1.

### Calculation of error decay rate

Given a matrix *A* that is the linearization of the discrete-time gradient descent dynamics in Equation 5, we compute the spectral radius as *ρ* = max{|*λ*| : *λ* is an eigenvalue of *A*}. Then the *error decay rate* is equal to the spectral radius *ρ*, and satisfies the bound *e*(*i*) ≤ *e*(0)*ρ*^*i*^ where *i* is iteration number.

## Data availability

All data are publicly available in a Code Ocean capsule, `DOI: 10.24433/CO.4049054.v3`

## Code availability

The data and analysis scripts needed to reproduce all figures and statistical results reported in both the main paper and supplement are publicly available in a Code Ocean capsule, `DOI: 10.24433/CO.4049054.v3`

## Acknowledgements

This work was supported by Meta Reality Labs Research, the National Science Foundation (NSF) award #2045014 (S.A.B.), the NSF award #2338662 (A.L.O) and the NSF award #2124608 (A.L.O and S.A.B.). M.M.M. was funded in part by the Department of Defense National Defense Science and Engineering Graduate Fellowship (NDSEG). The funders had no role in study design, data collection and analysis, decision to publish or preparation of the article. We thank A.X.T. Millevolte for experimental set-up support.

## Author Contributions

M.M.M., M.Y., S.A.B., and A.L.O., were responsible for experimental design. M.M.M, M.Y., and S.L. were responsible for experimental setup, M.M.M. and S.B. conducted the experiments, M.M.M. analyzed the experimental data and prepared figures, and M.M.M., S.A.B., and A.L.O. wrote the paper.

## Extended Data

### List of Extended Data Figures

**Extended Data Figure 1.**
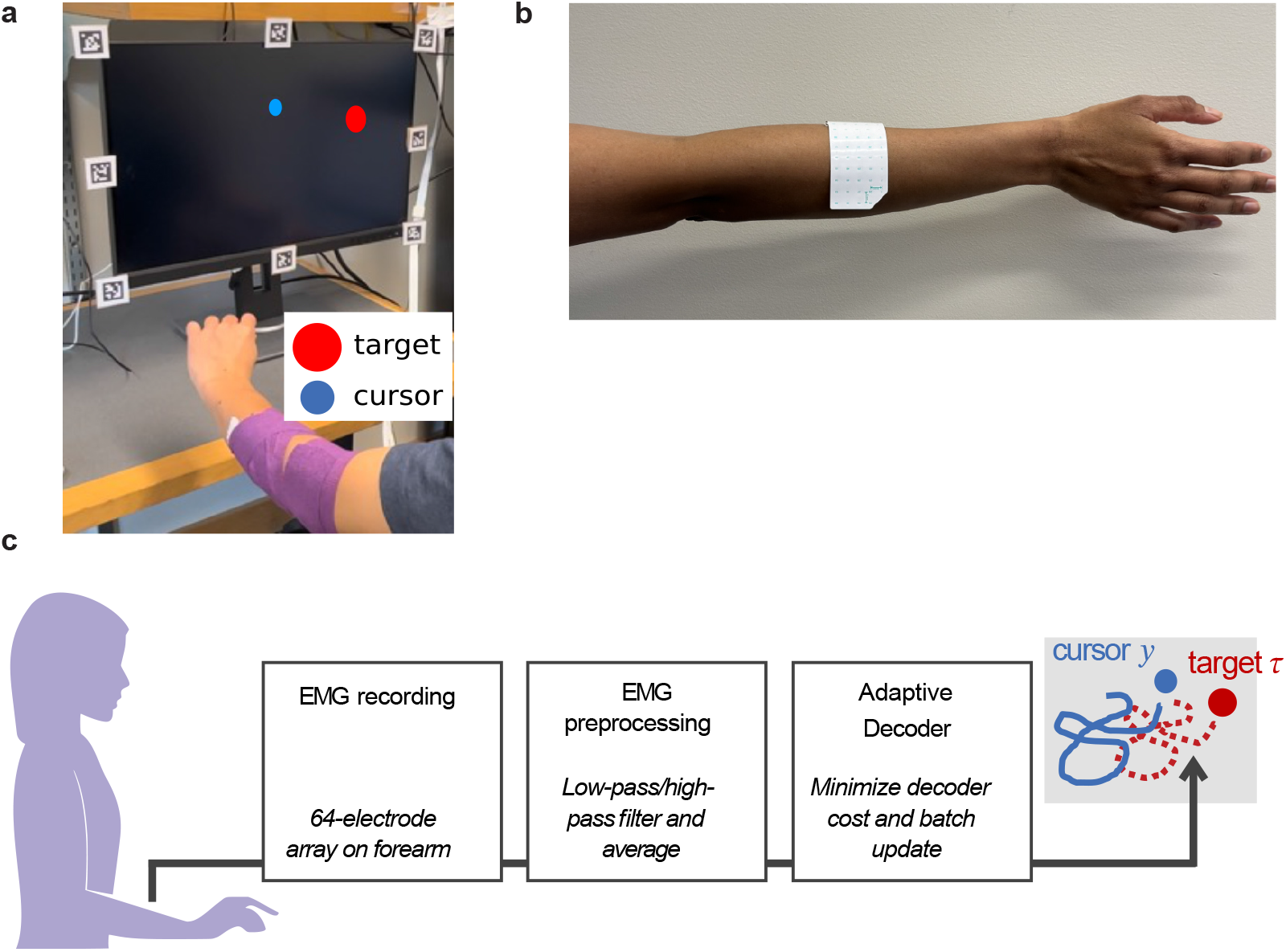
Experimental testbed. **a**. Photograph of experiment. User tracks target (dashed red) by controlling the velocity of a 2D cursor (solid blue) with their forearm muscle activity. Muscle activity is collected via surface EMG electrodes that are placed on the participant’s dominant forearm and wrapped with Coban tape. **b**. Electrode placement on dominant forearm. Surface EMG electrodes are placed on user dominant forearm to target Extensor Carpi Radialis. **c**. Schematic of EMG preprocessing and decoding pipeline. EMG signals recorded from the user forearm are preprocessed and then input to the adaptive decoder. The decoder output is the cursor velocity that is integrated to display cursor position (blue) on the screen.

**Extended Data Figure 2.**
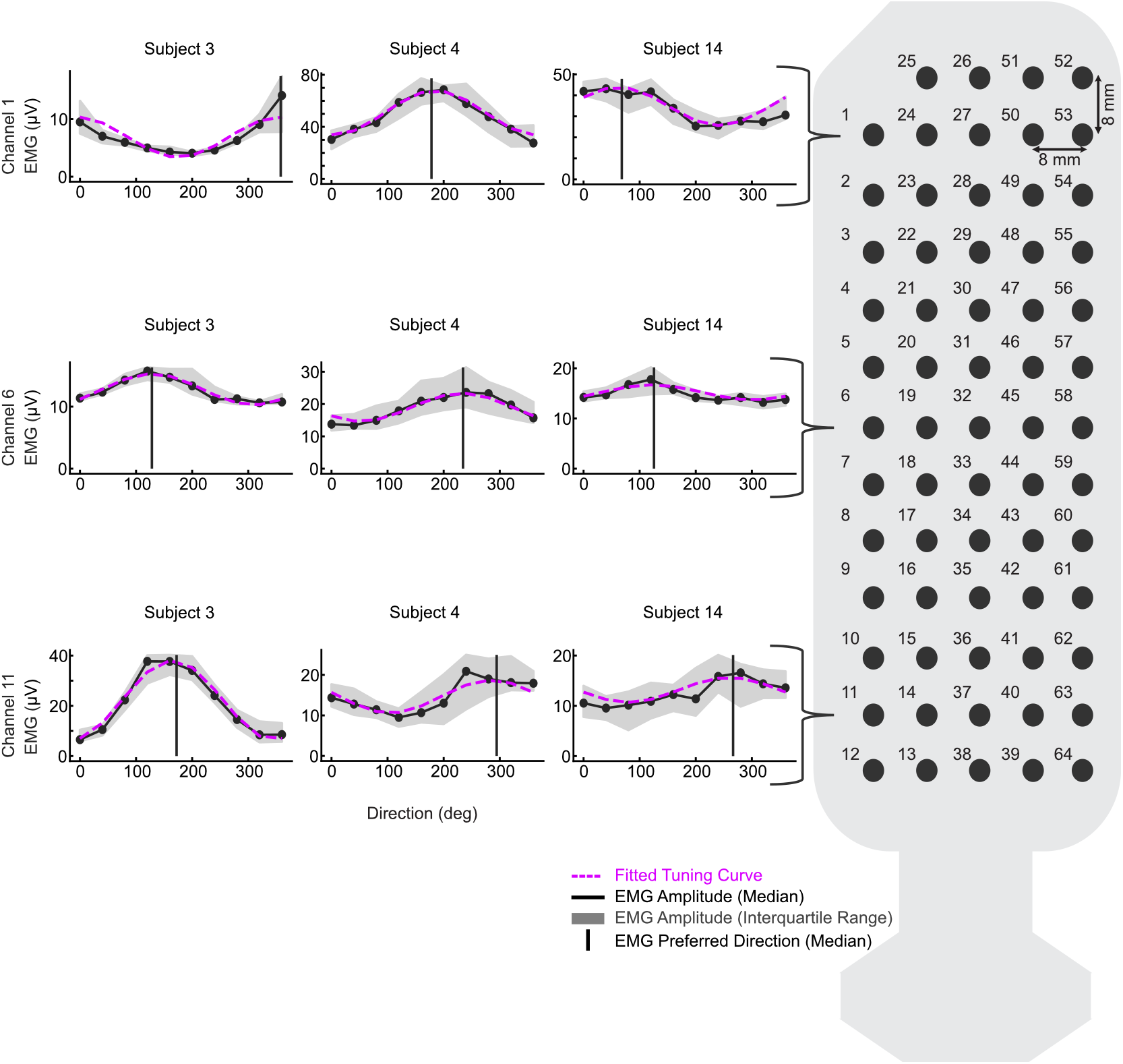
Surface EMG electrode and example tuning curves. Diagram of surface EMG electrode and sample EMG tuning curves from representative subjects taken from late segments within trial, black line represents median across trials with shaded area indicating the the 25th - 75th percentiles. Pink dashed lines show fitted tuning curves. Vertical lines represent preferred direction from late trial segments.

**Extended Data Figure 3.**
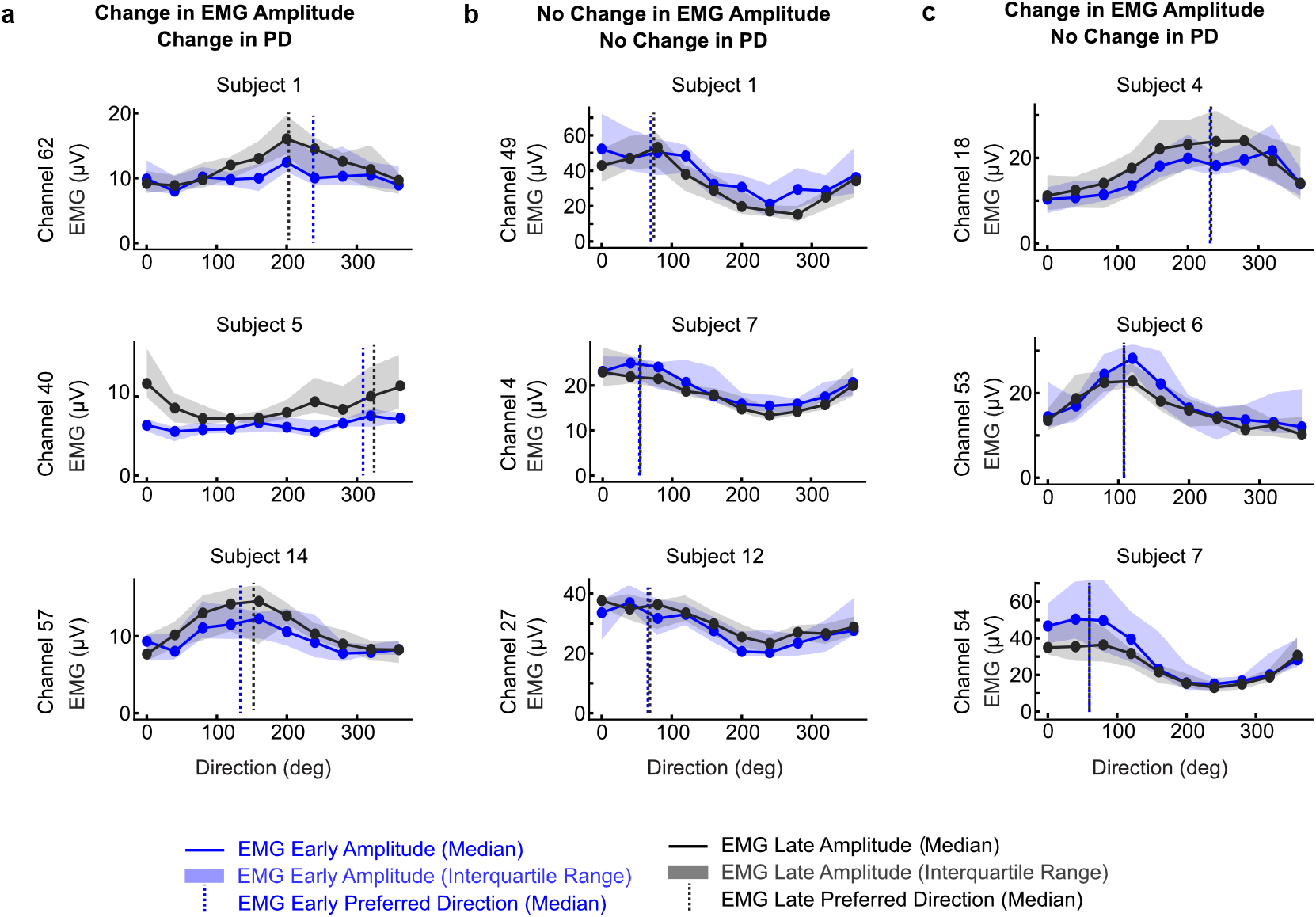
Example tuning curve changes from participants. Sample EMG tuning curves from range of participants taken from early (blue) and late (black) trial segments. For all plots, lines represent the median across trials with shaded area indicating the the 25th - 75th percentiles. Vertical dashed lines represent preferred direction (PD) from early (blue) and late (late) trial segments. Examples were chosen to illustrate the diversity of qualitative changes observed. **a**. Example EMG tuning curves showing qualitative differences in EMG amplitude and PD from early to late in the trial. **b**. Example EMG tuning curves showing minimal qualitative differences in EMG amplitude and PD from early to late in the trial. **c**. Example EMG tuning curves showing qualitative differences in EMG amplitude but minimal to no differences PD from early to late in the trial.

**Extended Table 1.**
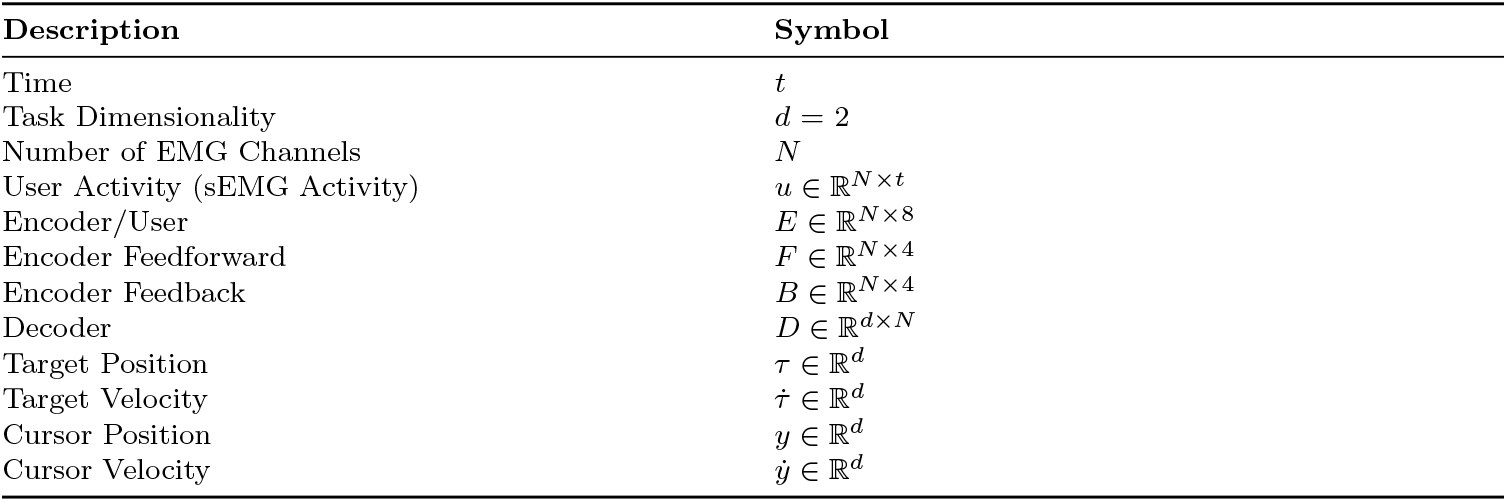
List of Model Variables.

| Description | Symbol |
| --- | --- |
| Time | $t$ |
| Task Dimensionality | $d = 2$ |
| Number of EMG Channels | $N$ |
| User Activity (sEMG Activity) | $u \in \mathbb{R}^{N \times t}$ |
| Encoder/User | $E \in \mathbb{R}^{N \times 8}$ |
| Encoder Feedforward | $F \in \mathbb{R}^{N \times 4}$ |
| Encoder Feedback | $B \in \mathbb{R}^{N \times 4}$ |
| Decoder | $D \in \mathbb{R}^{d \times N}$ |
| Target Position | $\tau \in \mathbb{R}^d$ |
| Target Velocity | $\dot{\tau} \in \mathbb{R}^d$ |
| Cursor Position | $y \in \mathbb{R}^d$ |
| Cursor Velocity | $\dot{y} \in \mathbb{R}^d$ |

**Extended Data Figure 4.**
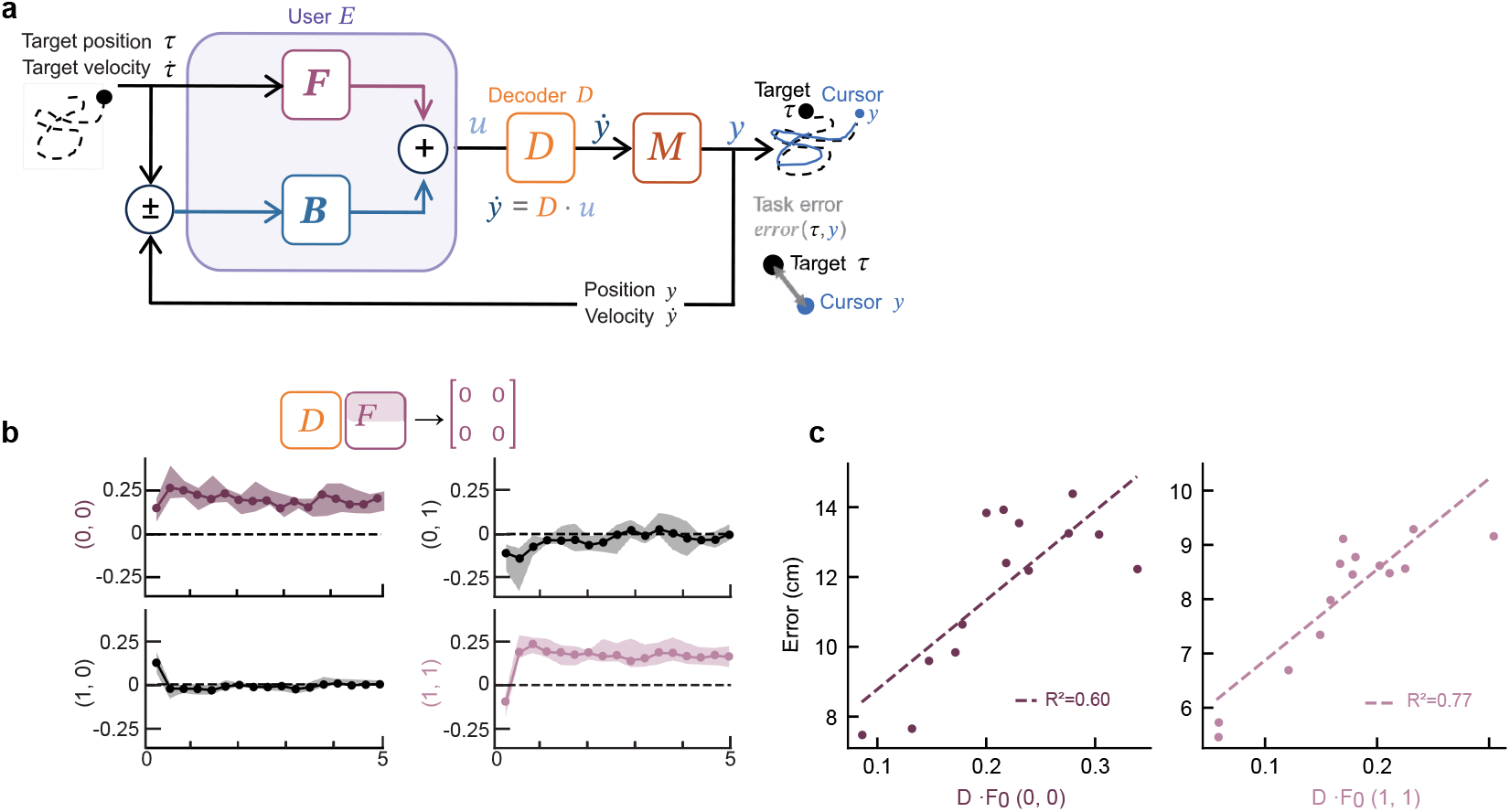
Effect of error on Decoder-Encoder Feedforward Pairs. **a**. Block diagram model of closed-loop system. The user’s encoder *E* outputs myoelectric activity *u* to the decoder *D* to follow the target’s position and velocity. The decoder outputs velocity, which is then integrated by system dynamics *M* to generate a cursor position. The user sees their cursor moving with some velocity 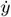 to position *y*. Error in the cursor position and velocity (relative to the target) acts as closed-loop feedback to the user. (Duplicated from Fig. 2A for completeness.) **b**. Product of the decoder matrix with first-order feedforward contributions of the encoder matrix (N = 14, median; shading shows the 25th - 75th percentile). Black dashed lines represent values that yield perfect trajectory tracking. Purple dashed lines represent values that yield near perfect trajectory tracking. (Duplicated from Fig. 2C, left for completeness.) **c**. Average tracking error (left: x-direction, right: y-direction) vs average *D* · *F*_0_ value for each individual participant (dot).

**Extended Data Figure 5.**
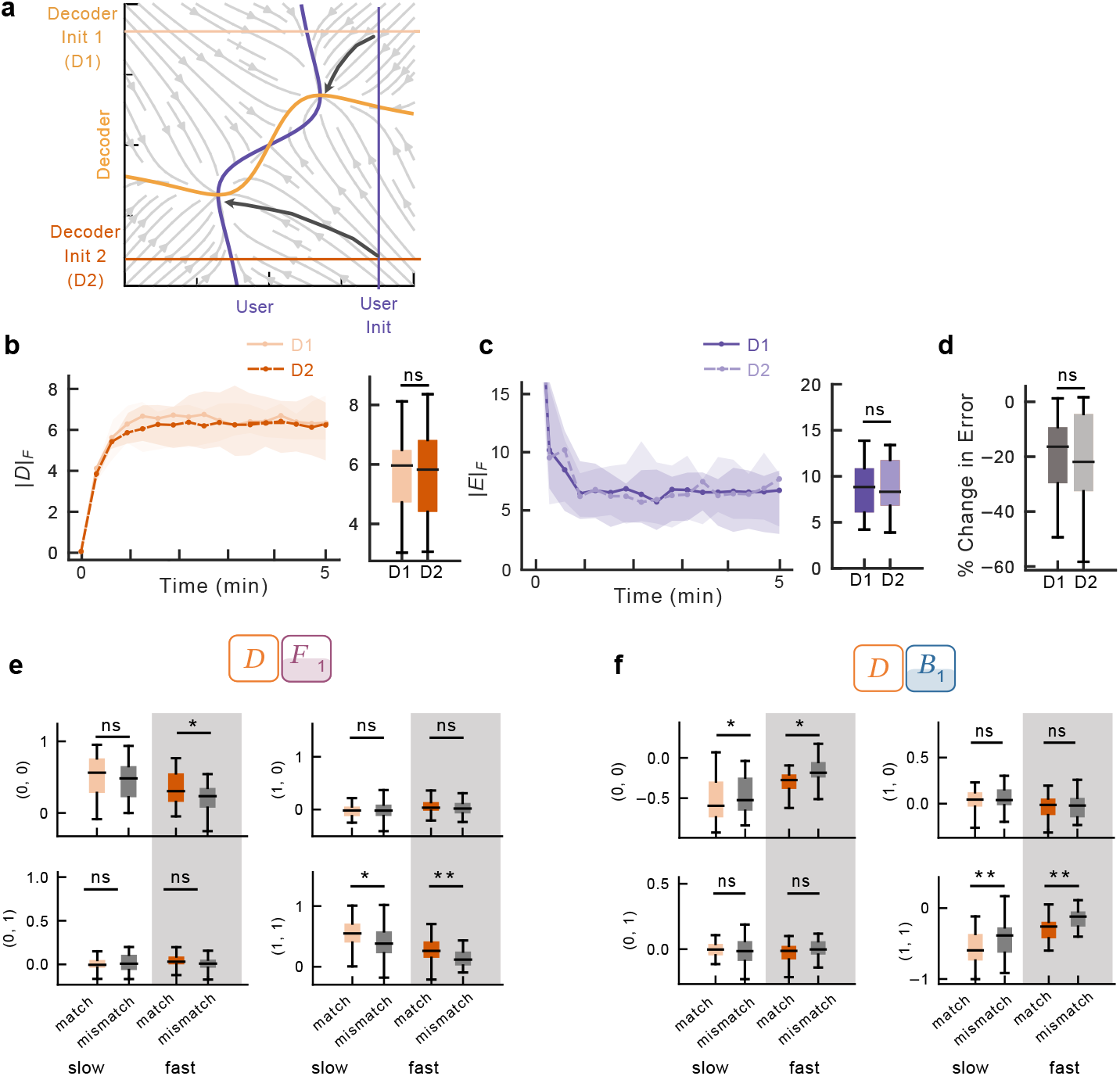
Decoder initialization slightly influences decoder-encoder pairs. **a**. Model predictions. All panels show the gradient field of the user and decoder cost functions. Purple (user) and orange (decoder) curves show nullclines (where the agent’s gradient equals 0) that intersect at stationary points (black stars). Decoder initialization D1 (light orange) and initialization D2 (dark orange) are noted on the vertical axis of the diagram. Assumed user initialization (purple) is noted on the horizontal axis. **b**. Left: Average magnitude of the decoder matrix (norm) as a function of time in the trial for D1 (solid light orange) and D2 (dashed dark orange) initializations in slow learning rate conditions (N = 14, median; shading shows the 25th - 75th percentile). Right: Boxplots are average decoder effort across the trial for each subject (N=14, center shows median; box shows 25th–75th percentiles; whiskers extend to 1.5 × this interquartile range; two-sided Wilcoxon signed-rank test, ns = 0.27). **c**. Left: Average magnitude of the user encoder matrix (norm) as a function of time in the trial for D1 (solid dark purple) and D2 (dashed light purple) decoder initializations in slow learning rate conditions (N = 14, median; shading shows the 25th - 75th percentile). Right: Boxplots are average effort for each subject across the trial (N = 14, center shows median; box shows 25th–75th percentiles; whiskers extend to 1.5 × this interquartile range; two-sided Wilcoxon signed-rank test, ns = 0.86). **d**. Percent change in error from the start of the trial (first 30 seconds) to the end of the trial (last 30 seconds), separated by D1 (dark gray) and D2 (light gray) initializations in slow learning rate conditions (N=14, center shows median; box shows 25th–75th percentiles; whiskers extend to 1.5 × this interquartile range; two-sided Wilcoxon signed-rank test, ns = 0.33). **e**. Last minute in trial of product of the average decoder matrix of each initialization with first-order feedforward (*F*_1_) contributions of the average encoder matrix of each initialization, separated by learning rate. Fast learning rate conditions are shaded gray. The matched conditions are the decoders and encoders of the same initialization and the mismatched conditions are the decoders and encoders of different initializations (N=28, center shows median; box shows 25th–75th percentiles; whiskers extend to 1.5 × this interquartile range; two-sided Wilcoxon signed-rank test, ns *>* 0.05; *p *<* 0.05; **p *<* 0.001). **f**. Last minute in trial of product of the average decoder matrix of each initialization with first-order feedback (*B*_1_) contributions of the average encoder matrix of each initialization, separated by learning rate. Fast learning rate conditions are shaded gray. The matched conditions are the decoders and encoders of the same initialization and the mismatched conditions are the decoders and encoders of different initializations (N=28, center shows median; box shows 25th–75th percentiles; whiskers extend to 1.5 × this interquartile range; two-sided Wilcoxon signed-rank test, ns *>* 0.05; *p *<* 0.05; **p *<* 0.001).

**Extended Data Figure 6.**
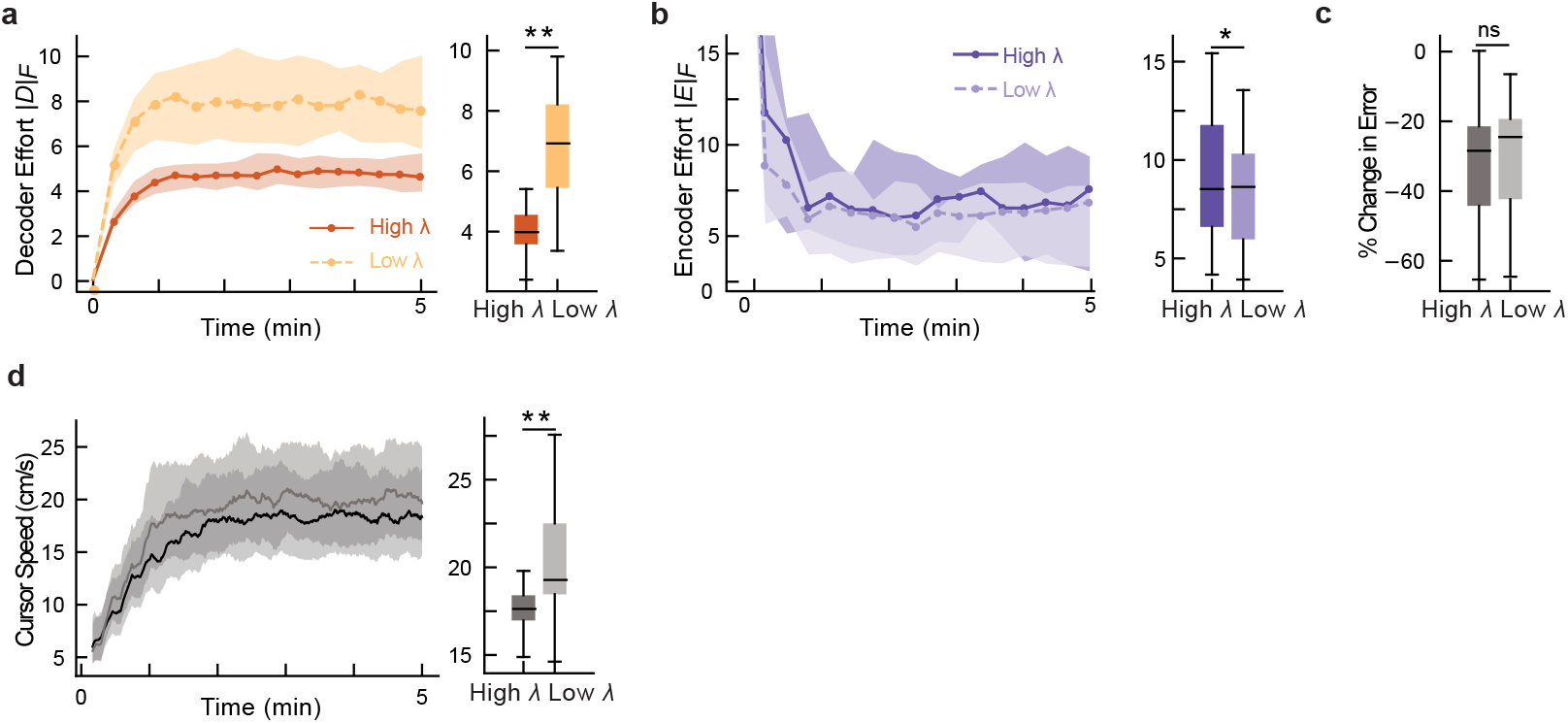
Effect of penalty parameter in slow learning rate condition only. **a**. Left: Average magnitude of the decoder matrix (norm) as a function of time in the trial for low (dashed light orange) and high (solid dark orange) decoder penalty terms (N = 14, median; shading shows the 25th - 75th percentile). Right: Boxplots are average decoder effort across the trial for each subject (N=14, center shows median; box shows 25th–75th percentiles; whiskers extend to 1.5 × this interquartile range ; one-sided Wilcoxon signed-rank test, **p = 6.1*e* − 5). **b**. Left: Average magnitude of the user encoder matrix (norm) as a function of time in the trial for low (dashed light purple) and high (solid dark purple) decoder penalty terms (N = 14, median; shading shows the 25th - 75th percentile). Right: Boxplots are average effort for each subject across the trial (N = 14, center shows median; box shows 25th–75th percentiles; whiskers extend to 1.5 × this interquartile range; one-sided Wilcoxon signed-rank test, *p = 0.018). **c**. Percent change in error from the start of the trial (first 30 seconds) to the end of the trial (last 30 seconds), separated by high (dark gray) and low (light gray) decoder penalty term conditions (N=14, median; shading shows the 25th - 75th percentile; two-sided Wilcoxon signed-rank test, ns = 0.24). **d**. Left: Average cursor speed as a function of time in the trial for low (light gray) and high (dark gray) decoder penalty terms (N = 14, median; shading shows the 25th - 75th percentile). Right: Boxplots are average cursor speed across the trial for each subject (N=14, center shows median; box shows 25th–75th percentiles; whiskers extend to 1.5 × this interquartile range; Wilcoxon signed-rank test, **p = 6.1*e* − 5).

## Footnotes

1 © 2022 Maneeshika M. Madduri, Momona Yamagami, Augusto X.T. Millevolte, Si Jia Li, Sasha N. Burckhardt, Samuel A. Burden, Amy L. Orsborn. Reproduced with permission from “Co-Adaptive Myoelectric Interface for Continuous Control.” IFAC PapersOnLine, 55/41 (2022), pp. 95-100. https://doi.org/10.1016/j.ifacol.2023.01.109

